# Homeostatic, repertoire and transcriptional relationships between colon T regulatory cell subsets

**DOI:** 10.1101/2023.05.17.541199

**Authors:** Deepshika Ramanan, Kaitavjeet Chowdhary, Serge M. Candéias, Martina Sassone-Corsi, Diane Mathis, Christophe Benoist

**Author notes:** Address correspondence to: Christophe Benoist, Department of Immunology Harvard Medical School, 77 Avenue Louis Pasteur, Boston, MA 02115. Salk Institute for Biological Studies, La Jolla, CA, USA.

## Abstract

Foxp3^+^ regulatory T cells (Tregs) in the colon are key to promoting peaceful co-existence with symbiotic microbes. Differentiated in either thymic or peripheral locations, and modulated by microbes and other cellular influencers, colonic Treg subsets have been identified through key transcription factors (TF; Helios, Rorg, Gata3, cMaf), but their inter-relationships are unclear. Applying a multimodal array of immunologic, genomic, and microbiological assays, we find more overlap than expected between populations. The key TFs play different roles, some essential for subset identity, others driving functional gene signatures. Functional divergence was clearest under challenge. Single-cell genomics revealed a spectrum of phenotypes between the Helios+ and Rorγ+ poles, different Treg-inducing bacteria inducing the same Treg phenotypes to varying degrees, not distinct populations. TCR clonotypes in monocolonized mice revealed that Helios+ and Rorγ+ Tregs are related, and cannot be uniquely equated to tTreg and pTreg. We propose that rather than the origin of their differentiation, tissue-specific cues dictate the spectrum of colonic Treg phenotypes.

## INTRODUCTION

The immune system in the digestive tract is exposed to a complex array of food antigens and microbes that provide essential support. Peaceful coexistence is essential, avoiding breach by microorganisms but also over-exuberant immune responses to their products that intrinsically tend to activate innate immune receptors. In addition to physical barriers (mucus layers, epithelial tight junctions) a state of “active tolerance” involves immunoregulatory circuits, among which are FoxP3^+^ T regulatory cells (Tregs). Tregs regulate mucosal immunity to both symbionts and pathobionts, acting on many types of immunocytes via anti-inflammatory cytokines and small molecule mediators, and help preserve intestinal physiology by promoting epithelial barrier functions and tissue repair ^1–4^.

Several subsets of intestinal Tregs have been described, characterized by their sensitivity to microbial influences, and by a particular transcription factor (TF), particularly Gata3, Helios, Rorγ and cMaf (reviewed in ^3, 4^). Schematically, Gata3^+^ Tregs mostly overlap with Helios^+^ Tregs, while cMaf^+^ Tregs largely overlap with Rorγ^+^ Tregs ^5–8^. For the latter, whether Rorγ or cMaf is the key TF driver is unclear ^7–10^. Rorγ^+^cMaf^+^ Tregs, but not Helios^+^Gata3^+^ Tregs, are strongly tuned by gut bacteria, rare in germ-free or antibiotic-treated mice, but are induced by a number of individual microbes belonging to different families ^7–11^. Helios^+^ Tregs are more abundant in the small intestine, while Rorγ^+^ Tregs dominate in the colon, plausibly tied to differences in microbial load in these locations. In addition, a sizeable population of Rorγ^-^Nrp1^-^ Tregs in the small intestine is modulated by dietary antigens ^12^. Importantly, activity of FoxP3, the key identifying TF for Treg cells, appears somewhat dispensable in Rorγ^+^ Tregs, while it is essential in Helios^+^ Treg cells ^13, 14^.

Treg cells can differentiate in the thymus, or in peripheral locations. It is often stated that Helios^+^Nrp1^+^ Tregs are “tTregs” that differentiated in the thymus, while Rorγ^+^ Tregs and related subsets are “pTregs” that derived from Tconv (conventional T) cells, under particular conditions of activation^8, 13, 15–17^, but it is unclear whether this correspondence is absolute ^4^.

Microbe-dependent Rorγ^+^ Tregs are thought to mediate tolerance to commensal and pathogenic bacteria. Rorγ^+^cMaf^+^ Tregs have been reported to dampen production of IFNγ, IL4 or IL17 by Teff (effector T) cells ^8–10, 18^, although different studies report different outcomes, perhaps resulting from variable microbial environments. Several studies suggest that deficiency in Rorγ^+^cMaf^+^ Tregs leads to more severe colitis ^7–10, 13, 16, 19^, intestinal mastocytosis, enhanced type 2 immune responses ^13, 20^, and susceptibility to food allergy ^21^. Seemingly contradictory effects of Rorγ^+^ Tregs on IgA production and coating of bacteria have been reported ^7, 18, 22, 23^. Noxious effects of Rorγ^+^ Tregs are also encountered in oncology: their frequency increases in colorectal cancer and they paradoxically promote IL17-mediated intestinal inflammation ^24, 25^. Helios^+^Gata3^+^ Tregs are thought to promote tissue repair, in good part because of their preferential expression of tissue repair transcripts (in particular *Areg*) ^5, 6, 26^. But Helios^+^Gata3^+^ Tregs also affect colitis in the T cell transfer model ^6^ and enteric graft-vs-host disease ^27, 28^. Helios^+^Gata3^+^ Tregs may also improve the outcome of colorectal cancer by suppressing Th17 responses and preventing excessive intestinal tissue damage ^29^. While these different roles have been proposed there have not been comparative studies of Treg subset functions.

The relative proportions of intestinal Treg subsets are tuned by a number of external influences. Several other cell-types influence intestinal Treg differentiation and homeostasis: CD103^+^ dendritic cells (DCs), CX3CR1^+^ macrophages, innate lymphoid cells (ILC) 3, eosinophils ^2^, and recently described populations of MHC-II^+^ Rorγ^+^ stromal/myeloid cells ^30–32^. The nervous system also influences the balance of intestinal Treg subsets ^33, 34^, involving substance-P and neuron-produced IL6 ^33, 35^, in a triangular crosstalk between gut microbiota, enteric or extrinsic neurons, and Tregs. The balance of intestinal Treg populations is under genetic control ^18^, with an intriguing maternal transmission of the setpoint of Rorγ^+^ Tregs that can carry across generations.

An integrated perspective on the identity and homeostatic regulation of intestinal Tregs is still lacking, as we have only a limited grasp of their population dynamics and inter-relationships. Do the different Treg subsets regulate each other, do they compete for the same homeostatic niches? How related and interconnected are they when compared by gene expression programs or by their TCR repertoires, and are their functions as distinct as previously reported? Further, it is unknown whether individual Rorγ^+^ Treg-inducing microbes elicit solely quantitative variations, or phenotypically different populations. For an integrative perspective, we analyzed in parallel a panel of mice with deficiencies in the four main driving TFs, which revealed imbricated genomic and functional programs controlled by each TF, while single-cell RNAseq and TCR sequencing showed that the main Treg subsets are more interconnected than previously thought.

## RESULTS

### Intestinal Treg subsets regulate each other to maintain a homeostatic balance

A variety of phenotypes are found among intestinal Treg cells, but their homeostatic relationships are unclear. As a means of introduction, and consistent with previous reports, the flow cytometry plots of colonic Treg cells of Fig. 1A lay out the major groups of intestinal Treg cells: Helios^+^ and Rorγ+ Tregs appear the most mutually exclusive, cMaf+ Tregs overlap mostly with Rorγ+ Tregs but some cMaf expression is also present in Rorγ-negative cells, and is detectable in Helios+ Tregs. Gata3 is expressed mostly by Helios+ Tregs, but again not exclusively. About 20% of colonic Tregs are Rorγ-Helios (hereafter double-negative, DN), only a minority of which express Gata3. FoxP3, the defining TF of Treg cells, was expressed largely identically in all subsets, perhaps slightly more in Helios+ Tregs (Fig. 1B). Colonic Rorγ+ Tregs appeared around weaning age ^9, 36, 37^. At postnatal day 5, almost all colonic Tregs were Helios+, DN Tregs expanding a week later, possibly tied to the introduction of solid food ^12^, followed by Rorγ+ Tregs, coinciding with the diversification of gut microbes (Fig. S1A).

**Figure 1:**
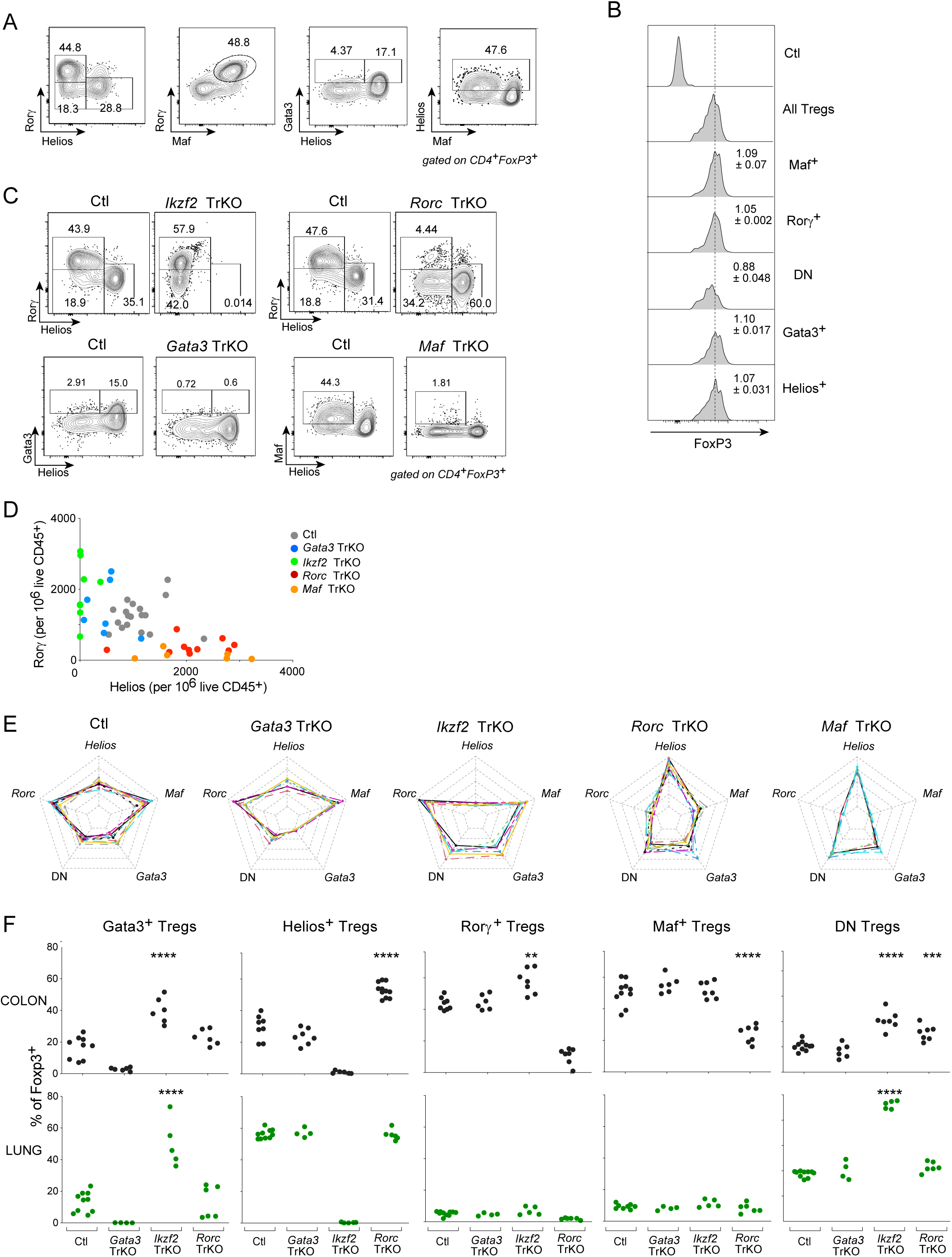
Homeostatic control of colonic Treg cells subsets by key transcription factors. (A) Representative flow cytometry plots of Treg subset defining transcription factors gated on TCRβ+ CD4+ Foxp3+ Tregs in wildtype SPF mice on the B6 background. (B) Representative histograms of Mean Fluorescence Intensity (MFI) of Foxp3 expression in different Treg subsets gated as in A. Values represent MFI normalized to all Tregs (Foxp3+), Ctl represents all Foxp3 CD4+ T cells. (C) Representative flow cytometry plots of intestinal Treg subsets in each Treg specific transcription factor knockout (TrKO) gated on TCRβ+ CD4+ Foxp3+ Tregs. (D) Quantification of total cell numbers of Rorγ+ Tregs vs Helios+ Tregs gated on TCRβ+ CD4+ Foxp3+ Tregs normalized to live CD45+ cells in each TrKO and littermate controls. Littermate controls for TrKOs were pooled together for comparison and all the mice were WT for the floxed allele and Foxp3-cre+. (E) Radar plots of total number of Treg cells expressing key transcription factors or DN (Rorγ Helios-) in each TrKO and pooled littermate controls. Limit of the radar plot was set from 0 to 100 and colors represent individual mice. (F) Proportions of different Treg subsets in colon (black) and lung (green) of each TrKO and pooled littermate controls. **p<0.01, and ****p<0.0001 by unpaired t-test.

To test comprehensively the homeostatic and functional relationships between these Tregs subsets, we generated a panel of conditional knockout mice with Treg-specific ablation of their defining TFs: Rorγ, Helios, Gata3 or cMaf (hereafter “TrKO”). In all these analyses, matching Cre-negative control littermates were included; as the results were essentially identical for all controls, we combined them here and in subsequent figures. Flow cytometric analysis confirmed loss of the TF (Fig. 1C), which was very complete, except for a small proportion of Tregs that remained Rorγ+ in *Rorc* TrKO mice (because the floxed Rorc allele is somewhat refractory to Cre-mediated deletion, and/or because of only recent expression of Foxp3-cre in pTregs). These Treg-specific ablations did not lead to visible pathology in the mice, but tuned the balance between various subsets in complex ways, as illustrated in Fig. 1D (cell numbers), and as a compilation of Treg proportions (Fig. 1E, F). Deficiencies in both Rorγ and cMaf led to marked decreases in Rorγ+ Tregs, and corresponding increases in Helios+ Tregs (Fig.1D). Conversely, Helios deficiency led to an increase in Rorγ+ Tregs (somewhat more modest, Fig. 1D) and in DN Tregs (Fig. 1F), the latter suggesting that some Helios+ Tregs persist in the absence of Helios, simply not recognizable as such. These results suggest that Helios+ and Rorγ+cMaf+ Tregs balance each other, likely by competing for the same homeostatic niche (or that they reciprocally inhibit each other). In contrast, the absence of Gata3 did not affect the proportions of any Treg subset (Fig.1D-F), suggesting that this TF does not control homeostatic regulation, rather effector functions. Unexpectedly, however, the absence of Helios led to an increase of Gata3+ Tregs (Fig. 1E, top left). Thus, even though Helios and Gata3 are expressed in many of the same Tregs, Helios appears to limit Gata3 expression (unless Gata3 increases to compensate for the missing Helios).

Since colonic Tregs can emigrate to extra-intestinal tissues {9441, 12647}, we also analyzed the effect of the same mutations in extra-intestinal tissues, such as the lung. In all the sites analyzed, the absence of Helios led to the same expansion of Gata3+ Tregs and DN Tregs as in the colon (Fig.1F bottom panel, S1B). On the other hand, the reciprocal increases of Helios+ and Rorγ+ Tregs in the absence of Rorγ/cMaf or Helios, respectively, were not observed outside the colon, likely because the microbial load and tissue environment are different, so the Rorγ+ Treg pool is small and unable to significantly replace Helios+ Tregs. These results suggest that the identifying TFs of colonic Treg subsets are differently required for the differentiation of the corresponding subsets, essential for Rorγ and cMaf, less so for Helios and Gata3.

### Differing functions of intestinal Treg subsets

As described above, specific functions in the control of intestinal inflammation have been ascribed to intestinal Treg subsets ^40^, but not in a comparative manner. We first performed a broad immunophenotyping of colon immunocytes in the TrKOs (salient results displayed in Fig. 2A, overall data aggregated in Fig. 2B). Numbers of lymphocytes (T, B or ILC) were not changed by any of the mutations. Consistent with previous reports ^8, 9, 18^, Th1 or Th17 Tconv cells (identified by production of IFNγ and IL17) were clearly elevated in both *Rorc* and *Maf* TrKO mice, while Th2 cells (identified by Gata3 expression) were actually reduced. *Ikzf2* TrKO yielded symmetrical trends in both respects (Fig. 2B). These observations are consistent with overlapping functions of Rorγ+ and cMaf+ Tregs, and with our prior results ^7–9, 18^, but differ somewhat from other reports ^7, 10^, possibly due to differences in microbiota across animal facilities. *Rorc* and *Maf* TrKO mice also showed an interesting decrease in Rorγ-expressing γδT cells, while total γδT cells were unaffected (Fig. 2A, B). This observation suggests a positive cooperativity between Rorγ+ Tregs and γδT cells; note that this does not extend to all Rorγ+ cells, as ILC3s, another dominant Rorγ+ cell-type in the intestine, were unchanged (Fig. 2A, B). The relationship between Rorγ^+^ Tregs and IgA production has been unclear ^7, 18, 22, 23^. In line with ref^7^, we found increased IgA+ plasma cells in colons of *Rorc* and *Maf* TrKO mice, most apparent with cMaf deficiency. (Total B cell numbers were unchanged.) Thus, overall, Rorγ+cMaf+ Tregs seem to have the strongest role in regulating the numbers and activation of other immunocytes in the gut.

**Figure 2:**
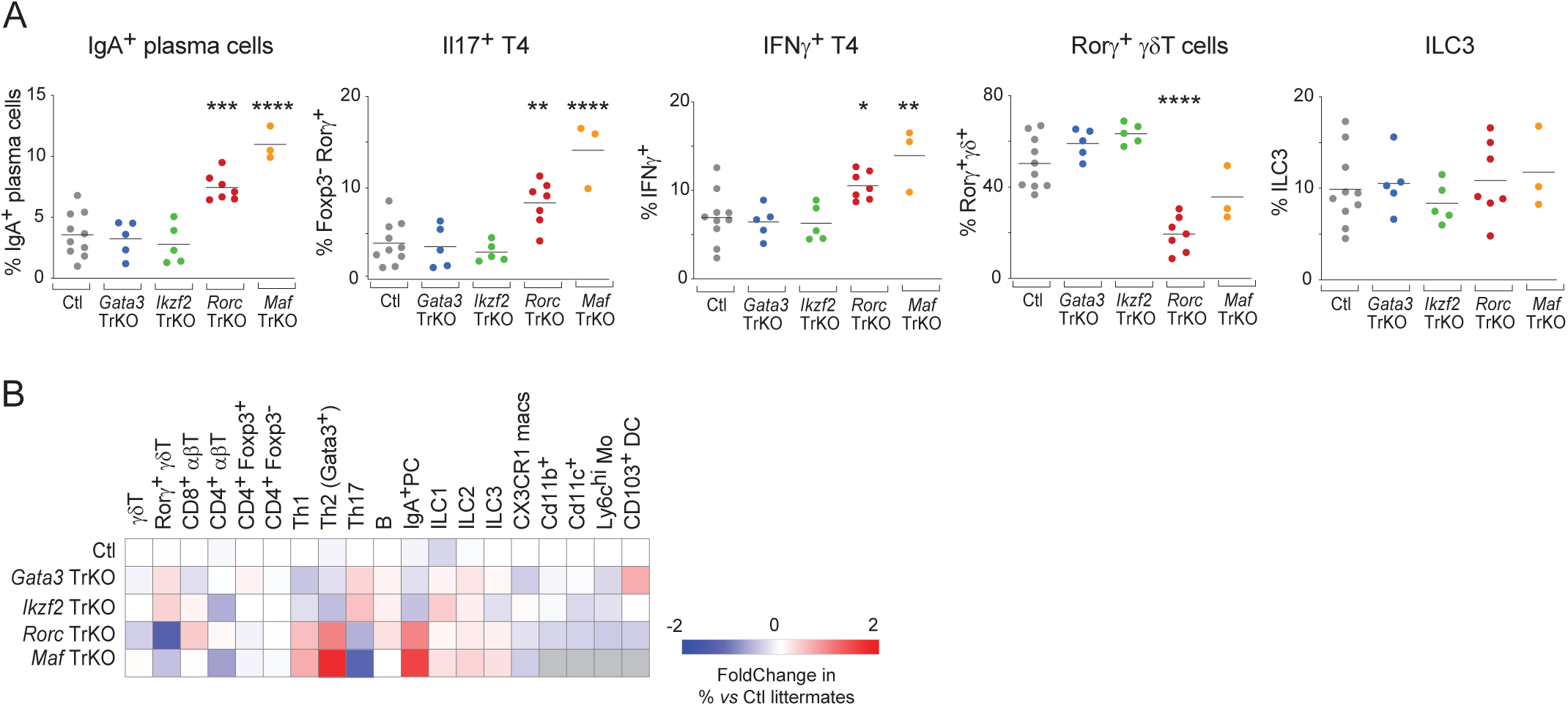
Rorγ+ cMaf+ Tregs regulate colonic lymphocyte populations. (A) Quantification of selected colonic immune cell types in different TrKO and pooled littermate controls. * p<0.05, **p<0.01, ***p<0.001, and ****p<0.0001 by unpaired t-test. (B) Heatmap of relative proportions of different immune cell types (normalized to WT littermate controls) in different TrKO mice.

We then tested the various conditional knockouts in infectious or inflammatory challenge models, studying the ensuing enteric response and inflammation. For these functional studies, wt/wt cre*+* littermates were used as controls, and all TrKOs were age-matched to within a week. First, we used a model of chemically induced colitis (dextran sodium sulfate – DSS – in the drinking water for 6 days + 4 days of recovery). *Rorc* TrKO mice developed more severe colitis (greater weight loss, shorter colons) compared with their littermate controls and other TrKOs (Fig.3A, B). DSS colitis normally entails a numeric increase in lamina propria Tregs. Here, this increase was observed with all TrKOs (Fig. 3C), with an increase in Rorγ+ Tregs (Fig.3D), but this did not happen in *Rorc* TrKO mice, suggesting that the adaptation to colonic inflammation predominantly involves Rorγ+ Tregs. The absence of Treg expansion in *Rorc* TrKO mice under DSS may well have contributed to greater inflammation (as might their increased Th1 and Th17 cells at baseline, which persisted under DSS Fig. S2A). Interestingly, while *Gata3* TrKO mice had a normal inflammatory response to DSS (Fig. 3A,B), particular adaptations were noted upon DSS challenge: an increase in CX3CR1+ macrophages, and a decrease in ILC2s post DSS (Fig.S2B), of unclear functional significance, but suggesting a specific role for Gata3+ Tregs in regulating the homeostasis of some immunocytes under challenge.

**Figure 3:**
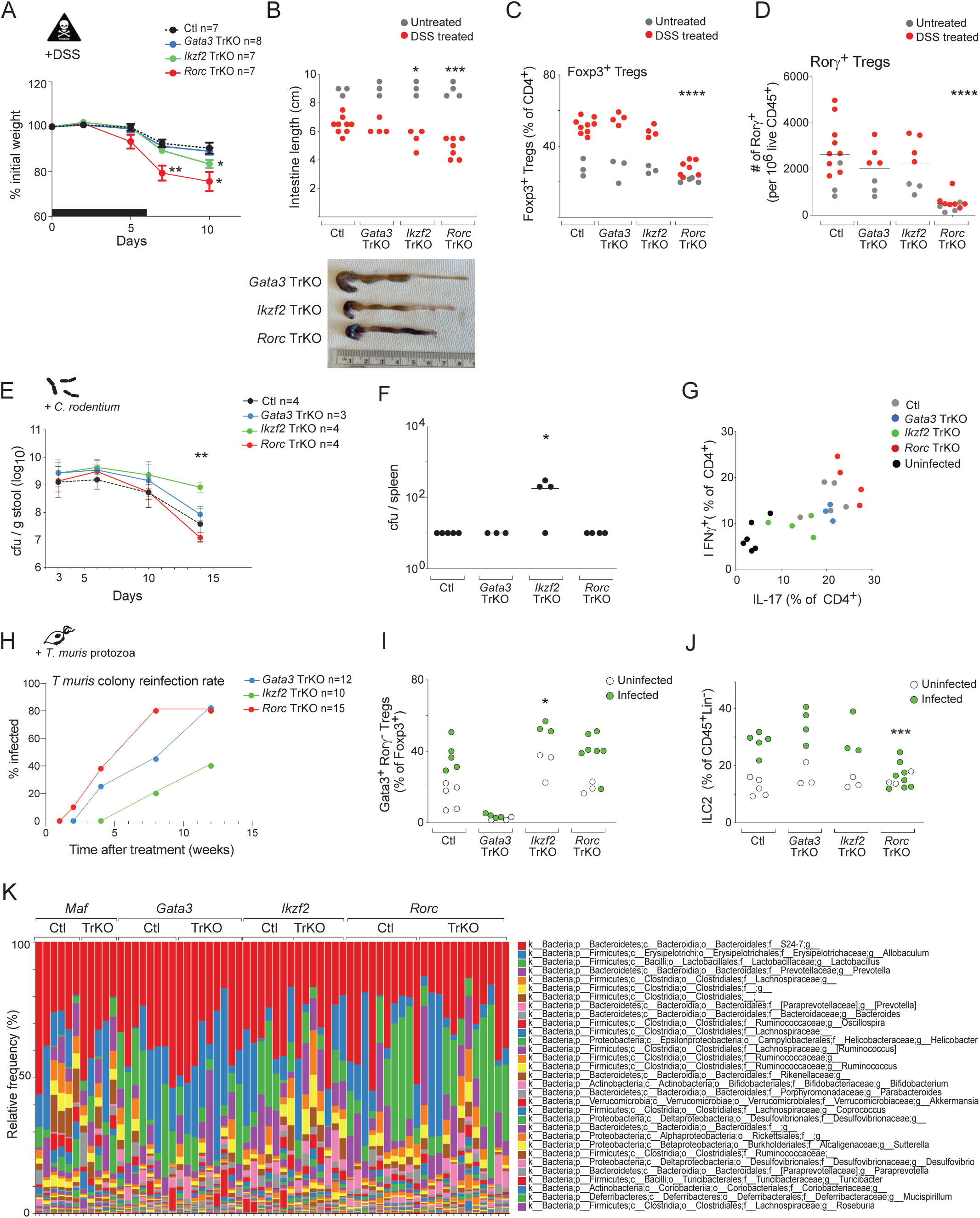
Regulatory functions of colonic Treg subsets are challenge dependent. (A) Survival curve of different TrKO mice and controls treated with 2.5% DSS at 6 weeks of age. * p<0.05, and **p<0.01 by unpaired t-test. (B) Quantification of colon length (measured from end of cecum to rectum) post-DSS treatment in untreated (gray) and DSS treated (red) TrKO mice and controls. * p<0.05, and ***p<0.001 by unpaired t-test (top). Representative picture of DSS treated colons from TrKO mice (bottom). (C) Proportions of Foxp3+ Tregs (gated on TCRβ+ CD4+) in untreated (gray) and DSS treated (red) TrKO mice and controls. ****p<0.0001 by unpaired t-test. (D) Proportions of Rorγ+ Tregs (gated on TCRβ+ CD4+ Foxp3+ Helios-) in untreated (gray) and DSS treated (red) TrKO mice and controls (top). Quantification of total cell numbers of Rorγ+ Tregs normalized to live CD45+ cells in TrKO and controls. ****p<0.0001 by unpaired t-test. (E) *C. rodentium* cfu recovered from stool over the course of infection in TrKO mice and controls. **p<0.01 by unpaired t-test. (F) *C. rodentium* cfu recovered from spleen on day 15 post-infection in TrKO mice and controls. *p<0.05 by Mann-Whitney test. (G) Representative flow cytometry plots of IFNγ vs IL-17 in TCRβ+ CD4+ Foxp3 T cells in the colon of *C. rodentium* infected TrKO mice and their pooled littermate controls and uninfected control mice. (H) Quantification of *T. muris* reinfection rate (% of mice with resurgence of *T.muris* post metronidazole treatment) in TrKO mice. (I) Proportions of Gata3+ Rorγ− Tregs in TrKO mice highly infected with *T.muris* (see methods for low vs high infection determination). *p<0.05 by unpaired t-test. (J) Proportions of Gata3+ ILC2 (% CD45+ Cd11b Cd11c B220 Ter119 TCRβ TCRδ NK1.1-) in TrKO mice highly infected with *T.muris.* ***p<0.001 by unpaired t-test. (K) Relative abundance of bacterial genus in stool of TrKO mice of the indicated groups generated by 16S rRNA sequencing.

We also tested the response of the TrKO mouse panel during gastrointestinal infection with *Citrobacter rodentium.* We monitored bacterial colonization (cfu), weight loss, diarrhea, intestinal pathology, and analyzed various immune cell populations, including the expected Th17 response. Here, it was the *Ikzf2* TrKO mice that stood out, with an increased bacterial burden relative to control littermates and other TrKOs, and bacterial dissemination to systemic organs, here the spleen (Fig.3E, F). *Ikzf2* TrKO mice also had a reduced Tconv response (IL17 and IFNγ) to *C. rodentium* (Fig.3G). These differences suggest a paradoxical role of Helios+ Tregs in favoring responses to this pathogen, although the effects might be due to the compensatory increase, shown above, in Rorγ+ Tregs in the absence of Helios+ Tregs (Rorγ+ Tregs acting as dominant suppressors of the *Citrobacter*-specific response).

Finally, we analyzed responses to a common protozoan parasite, *Tritrichomonas muris*, a Th2 inducing parasite ^41^, which is endemic in certain SPF facilities since it is not a specific pathogen that mice are routinely tested for. In order to remove the contamination, we treated the TrKO lines with metronidazole, a common treatment for *T. muris*. Because the parasite is easily transmissible, the lines became re-infected over time, but we noticed on several occasions (3 times over a 12 month period) that *Rorc* TrKO cages were most susceptible to *T. muris* reinfection, as illustrated for one time period during which reinfection was actively tracked (Fig.3H). We used this opportunity to investigate the influence of Treg subsets on other immunocytes during *T. muris* infection (considering only mice with a high parasite burden). Gata3+ Tregs were consistently increased in control mice and across the different TrKOs (Fig.3I; other Treg subsets being unchanged). This increase in Gata3+ Tregs in *Ikzf2* TrKOs further supported the notion that Helios is dispensable for the existence of Helios+Gata3+ Tregs. We also noted increases in ILC2s upon *T. muris* infection, as might be expected ^41^; this response appeared normal in *Gata3* and *Ikzf2* TrKOs, but was curtailed in *Rorc* TrKOs (Fig. 3J). These complex results (dominant expansion of Gata3+ Tregs, ineffective anti-protozoan response in *Rorc* TrKOs) suggest an interplay between intestinal Treg subsets. They differ from previously reported results with the helminth *Heligmosomoides polygyrus*, against which responses were more effective in Rorγ+ Treg-deficient mice ^10^. These differing outcomes could be due to differences in the parasite (helminth vs protozoan), or in resident microbiota (given that Ohnmacht et al noted increased Th2 in their *Rorc* TrKO mice).

As another test of differential function of intestinal Treg subsets, we assessed whether any of the TrKO lines harbored changes in their intestinal bacterial content, by performing 16S rDNA profiling of stool bacteria. Given the known issues with cage-of-origin fluctuations in microbiota, our protocol compared pairs (each TrKOs had one matched control littermate; n= 6 to 12 TrKOs per line). As illustrated in Fig. 3K, S3A, and as confirmed by the rarefaction analysis of Fig. S3B, there was no strong dysbiosis in any of the TrKO lines. We detected only a few differentially represented Amplicon Sequence Variants by QIIME2 (Fig. S3C), in numbers comparable with noise estimated by permutation. Overall, these comparative analyses of host-microbe interactions indicated that the Treg subsets work in concert to maintain different aspects of intestinal immune function in the context of microbial challenges.

### Genomic Identity of Intestinal Tregs

The results above indicate that these key TFs have different effects, either conditioning the existence of intestinal Treg subsets or modulating their functional capabilities. It was important, then, to determine their impact on gene expression programs, as inscribed in the genome by chromatin accessibility patterns. To systematically examine the impact of each TF on genome-wide chromatin states, we performed single cell assay for transposase-accessible chromatin (scATAC-seq) on colon Tregs from each of the four TrKO lines and littermate controls (each in duplicate). Gene scores, chromatin-based proxies for gene expression ^42^, clearly delineated Helios+ and Rorγ+ Treg populations on a 2D UMAP (Uniform Manifold Approximation and Projection) visualization of the data (Fig. 4A). Relative accessibility of an ATAC-seq signature of Helios vs Rorγ Tregs from a prior study ^14^ supported these annotations (Fig. 4B). In accordance with flow cytometry results (Fig. 1B), *Foxp3* gene scores were similarly distributed across all Tregs (Fig. 4A), whereas *Gata3* predicted expression was restricted to the Helios+ population and *Maf* activity spanned Rorγ+, a subset of Helios+, and DN populations.

**Figure 4.**
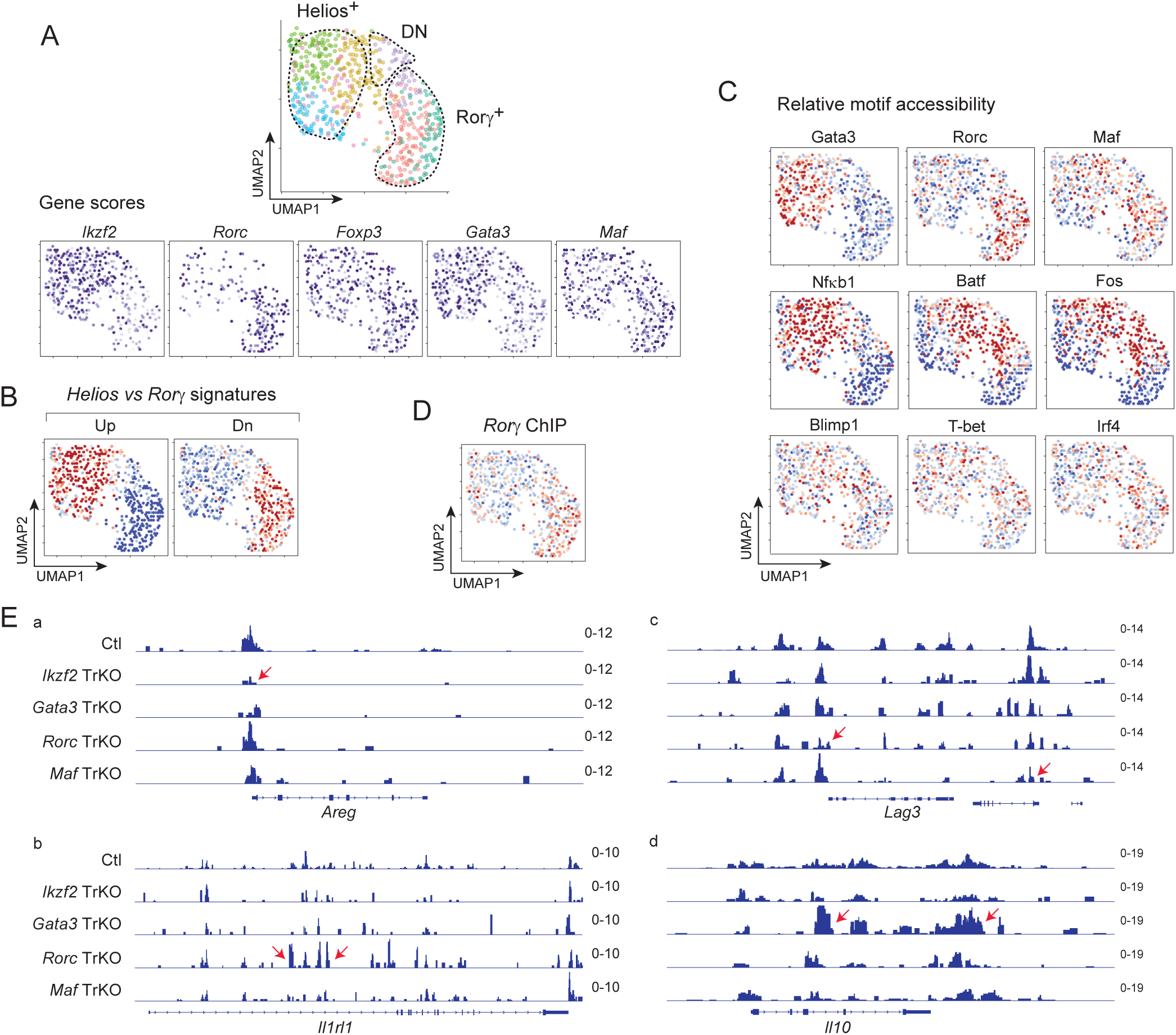
Specific genomic identities within the colonic Treg space. (A) UMAP of scATAC-seq data, grouping cells from all mice (TrKO and WT littermates). Gene scores (chromatin-based proxy for gene expression) for select genes are overlaid onto the UMAP. (B) Relative per-cell accessibility (chromVAR scores) of OCRs that distinguish Helios and Rorγ populations (from ^14^), overlaid onto UMAP from A. (C) Relative per-cell accessibility (chromVAR scores) of OCRs containing target motifs for the indicated transcription factors, overlaid onto UMAP from A. (D) Relative per-cell accessibility (chromVAR scores) of OCRs within genomic regions bound by Rorg (from ^44^), overlaid onto UMAP from A. (E) Aggregated accessibility profiles from the scATAC-seq data at the Areg, Il1rl1, Lag3, and Il10 loci for Tregs from Ctrl or TrKO mice. Arrows indicate regions with genotype-specific loss or gain of accessibility signal.

We next examined the activity of TF target sites across the cell populations by computing the per cell relative accessibility of open chromatin regions containing each TF motif (Fig. 4C). This mode of analysis reveals the cells in which each TF actually impacts the chromatin accessibility of its target sites ^14, 43^. Consistent with the gene scores, Gata3 motif accessibility was highest in the Helios+ Treg population, well demarcated from Rorγ motif accessibility (Fig. 4C, top row). The latter was confirmed by the accessibility distribution of known Rorγ-bound sites determined by chromatin immunoprecipitation ^44^ (Fig. 4D). Helios+ Tregs also had higher accessibility of NF-κB motifs, consistent with previous results identifying the activation of TNFRSF-NF-κB related pathways among ST2+ Tregs ^45, 46^. On the other hand, other motifs also had overlapping patterns of accessibility across the Treg space, reflecting the combinatorial and interwoven nature of TF operation in Tregs ^14^: Maf motif-containing regions had high accessibility in most Rorγ+ Treg cells but also a segment of Helios+ Tregs, in accordance with the flow cytometry data of Fig. 1; accessibility associated with BATF and AP-1 (e.g., Fos) motifs cut across Helios+ and Rorγ+ Tregs, through DN Tregs (Fig. 4C). Other TFs that have been associated with distinct functional subsets of Tregs (T-bet, IRF4, and Blimp-1 motifs) had even more diffuse activity (Fig. 4C, bottom row). Thus, each subpopulation of Tregs had distinct TF target activities.

Overlaying the distribution of cells from each TrKO genotype onto the UMAP space (Fig. S4) brought a pattern consistent with the flow cytometry results of Fig. 1. Whereas WT cells were represented in Helios+, Rorγ+, and DN Treg populations, *Ikzf2* TrKO and *Rorc* TrKO Tregs were shifted to Rorγ+ and Helios+ populations respectively. Echoing the heterogenous expression and effects of cMaf, *Maf* TrKO Tregs were shifted to both the Helios+ and DN populations and away from the Rorγ+ region.

Examining aggregated scATACseq reads at regulatory regions around Treg effector genes revealed varied regulatory strategies (Fig. 4E). For *Areg*, a locus known to be over-expressed in Helios+Gata3+ Tregs, the major effect was a reduction of accessibility around its promoter region in the *Ikzf2* TrKO (Fig. 4E.a). On the other hand, around the co-expressed *Il1rl1* locus (encodes the IL33 receptor), the dominant effect was an increase in signal in the *Rorc* TrKO (Fig. 4E.b), possibly resulting from release of Rorγ mediated repression. (Note that this effect is not observed in the *Maf* TrKO.) For effector molecules generally associated with the Rorγ+Maf+ Treg contingent, reduced accessibility was observed at several regulatory elements within the *Lag3* gene body in *Rorc* and *Maf* TrKO Tregs (Fig. 4E.c), suggesting a positive control by both factors. (Interestingly, the lost signal was not identical for both.) On the other hand, accessibility at regulatory regions of the *Il10* locus was strikingly boosted in *Gata3* TrKO Tregs (Fig. 4E.d), perhaps again reflecting a de-repression event. Thus, each TrKO also had consequences for the regulation of molecules important for Treg effector function, providing some molecular basis for the divergence of phenotypes following immunologic challenge, and again highlighting different regulatory strategies used by these TFs.

### How individual microbes mold intestinal Tregs

Individual microbes can induce intestinal Tregs, particularly Rorγ+ Tregs, in different proportions; several different mechanisms have been invoked to explain these differences ^1, 2, 4, 40^. These inductive events are observed as an increase in Treg proportions; yet it is unclear what underlying cellular changes are actually involved (accelerated pTreg conversion, Treg expansion, reduced cell death, etc)^4^. It is also unknown whether the Tregs amplified by different microbes are phenotypically the same, and whether recognition of antigenic epitopes on inducing microbes is at play. To address these questions, we performed single cell RNA sequencing (scRNAseq) on activated CD4+ T cells (TCRβ+CD4+CD44hi) from the colons of germ-free mice (GF) mono-colonized with individual microbes. We selected a high inducer of Rorγ+ Tregs (*Clostridium ramosum)*, an intermediate inducer (*E. coli Nissle)* and *Peptostreptococcus magnus*, which is unable to induce Treg cells ^47^, also including SPF B6 mice as controls. The single cell profiling was performed three independent times, each with biological replicates of each condition, multiplexed together with DNA “hashtags” to ensure optimal comparability (a total of 24 mice; results from one experiment are presented in the main figures, replicate in Fig. S5B). Dimensionality reduction and visualization showed that all mice displayed the expected populations (based on gene signatures): naive CD4 T cells, conventional/effector T cells, and Tregs (Fig. S5A). We then focused the analysis on Treg cells, which could readily be classified into 3 clusters, Helios+, Rorγ+ and DN Tregs based on previously reported signatures ^48^ (Fig. 5A). As expected, GF+ *C. ramosum* and GF+ *E. coli Nissle* had more Rorγ+ Tregs compared with GF mice or GF+ *P.magnus* (Fig. 5B). The distribution of cells across the gene expression space suggested that the differences between Treg populations in the presence of the different microbes were mainly quantitative. Colon Tregs from mice colonized with *C. ramosum* simply had a more skewed distribution towards the Rorγ+ Treg quadrant (Fig. 5B). Differential Gene Expression analysis, factoring out these numerically skewed distributions, uncovered no microbe-specific geneset. This conclusion was confirmed by plotting differentially expressed genes: while there were clear differences between Treg subsets, none was microbe specific (Fig. 5C). Thus, whatever means *C.ramosum* and *E. coli* may use to induce Rorγ, they do so by titrating more or less of the same cell, not different cell-types.

**Figure 5.**
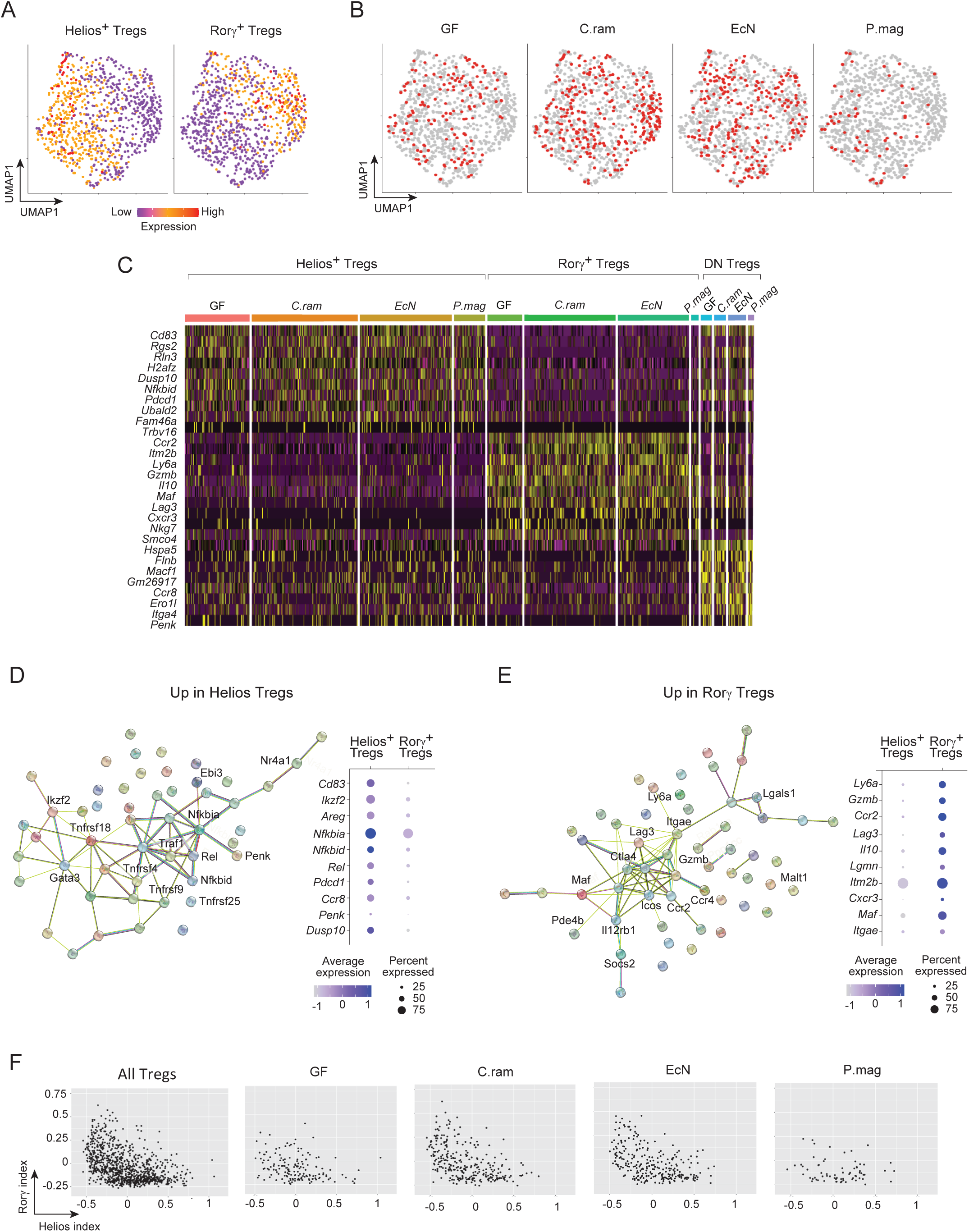
Helios+ Tregs and Rorγ+ Tregs have unique transcriptional signatures that are conserved across responses to individual microbes. (A) UMAP projection of colonic Treg cells from a representative scRNAseq analysis of colonic Tregs in GF (germ-free) and monocolonized mice, color-coded for expression of RORγ+ and Helios+ Treg gene signatures (genes in Materials and Methods). Repeat in Fig. S5. (B) UMAP projection of colonic Treg cells as in (A), split between hash tagged cells from different mice. Each hashtag represents an individual mouse: GF control, and GF mice monocolonized with C.ram (*C. ramosum*), EcN (*E. coli* Nissle), and P.mag (*P. magnus*). (C) Heatmap of top differentially expressed genes from each cluster (Helios+, Rorγ+ or DN) across the different Treg subsets from GF or monocolonized mice. (D) String analysis and dotplot of top differentially expressed genes in Helios+ Tregs pooled from all GF and monocolonized mice. (E) String analysis and dotplot of top differentially expressed genes in Rorγ+ Tregs pooled from all GF and monocolonized mice. (F) 2D representation of Tregs based on gene-signature scores calculated for Helios+ Tregs and Rorγ+ Tregs (genes used to calculate scores in Materials and Methods).

We then used the dataset to generate a more refined transcriptional signature for Helios+ and Rorγ+ Tregs (Table S1, Fig. 5C-E). The results extended and refined our prior signatures ^9, 48^. Helios+ Tregs were marked by increased expression of some expected genes such as *Gata3*, *Cd83*, *Il1rl1* and *Areg*, but they also displayed an increase in NF-κB signaling-related genes, in keeping with the stronger accessibility of NF-κB motifs observed in the scATACseq data above. Rorγ+ Tregs displayed increased *Maf*, *Ccr2* and *Lag3*, but they were also marked by increased effector molecules like *Gzmb* and *Il10*, as also expected from the results above. Interestingly, the DN Treg population was characterized by stronger expression of a small geneset, including the neurotransmitter *Penk* and *Ccr8*, a molecule of interest in tumor Tregs.

To ask if this refined signature might help sharply distinguish colonic Treg subsets, we calculated Helios+ Tregs and Rorγ+ indices for each cell, and visualized the data on a 2D plot (Fig. 5F). Even with refined signatures, the scRNAseq profiling indicated that the demarcation between Treg subsets was not sharp, similar to what was observed between Helios+, Rorγ+ and DN subsets by flow cytometry. Both genomic and protein profiling suggest that while there are distinct Treg subsets, they exist in a continuum that is modulated by microbes and the local environment, rather than as discrete and distinct cell types.

### TCR selection, diversity and clonality in response to defined intestinal microbes

These scRNAseq data allowed us to relate the differentiated phenotypes of Treg cells in the monocolonized mice with the composition of the TCRs expressed by each cell. The nucleotide sequences of rearranged clonotypes can serve as molecular barcodes to identified clonally related T cells, which here should help elucidate differentiation paths of the different Treg subsets. From three independent profiling experiments, we obtained the full sequence of TCRα/TCRβ pairs for 24,366 colonic CD4+ T cells from 24 GF, monocolonized (*C. ramosum* or *E. coli*), or SPF mice (with matched spleen data for 10 of these mice; Table S2A/B). In line with our previous report ^49^, the T cell repertoire in monocolonized mice was diverse, with a broad usage of *TRAV* and *TRBV* regions (Table S2C).

We first analyzed these sequences for clues to the timing of differentiation of these cells, given that Treg cells generated in early periods have particular properties ^50–52^. TCR rearrangements in pre-T cells during the fetal and perinatal periods lack N region diversity, because of ontogenically delayed expression of TdT ^53, 54^. Examination of *TRA* and *TRB* rearrangements showed that the frequency and size of N nucleotide additions, as well as base deletions from the recombining ends, were similar in colonic Helios+ and Rorγ+ Tregs, as well as colonic Tconv, and in the same range when compared to splenic T cells from SPF mice (Table S2D). Furthermore, rearrangements at the *Tcra* locus are not completely uniform across ontogenic time, with a propensity for the more proximal *Va* and *Ja* genes to recombine in T cells generated in the fetal and perinatal times ^55, 56^. *Tcra* sequences from Rorγ+ and Helios+ Tregs showed a broad distribution of *TRAV* and *TRAJ* usage, comparable with that of Tconv cells in the colon, with no preference for joins involving proximal Va regions (Fig. S6). These results suggest that neither Helios+ nor Rorγ+ Tregs are enriched in cells that differentiated during the fetal/peri-natal period.

As previously noted, repertoires of colonic CD4+ T cells from monocolonized mice were diverse, but also included a notable proportion of clonotypes present in several cells (Fig. 6A/B). These repeated clonotypes stemmed from clonal expansion from a common parent cell, because their nucleotide sequences were completely identical at both *Tcra* and *Tcrb* loci in all cells expressing that clonotype in a given mouse, with frequent N nucleotide addition indicating that they did not derive from favored homology-driven rearrangements (Fig. S7). In contrast, in some instances where the same protein clonotypes occurred in different mice, they were always encoded by different nucleotide sequences (Fig. S8, showing that identical TCRα and TCRβ rearrangements in Tregs of different phenotypes did not result from favored recombination events. These expanded clonotypes were represented at similar frequencies in colonic Treg and CD44^hi^ Tconv cells, but were less frequent in Tconv cells with naïve phenotypes, indicating that they were triggered by antigen exposure. Interestingly and perhaps unexpectedly, monocolonization did not lead to narrower expansions than seen in colons of SPF mice, as evidenced by the rarefaction plots of Fig. 6B.

**Figure 6.**
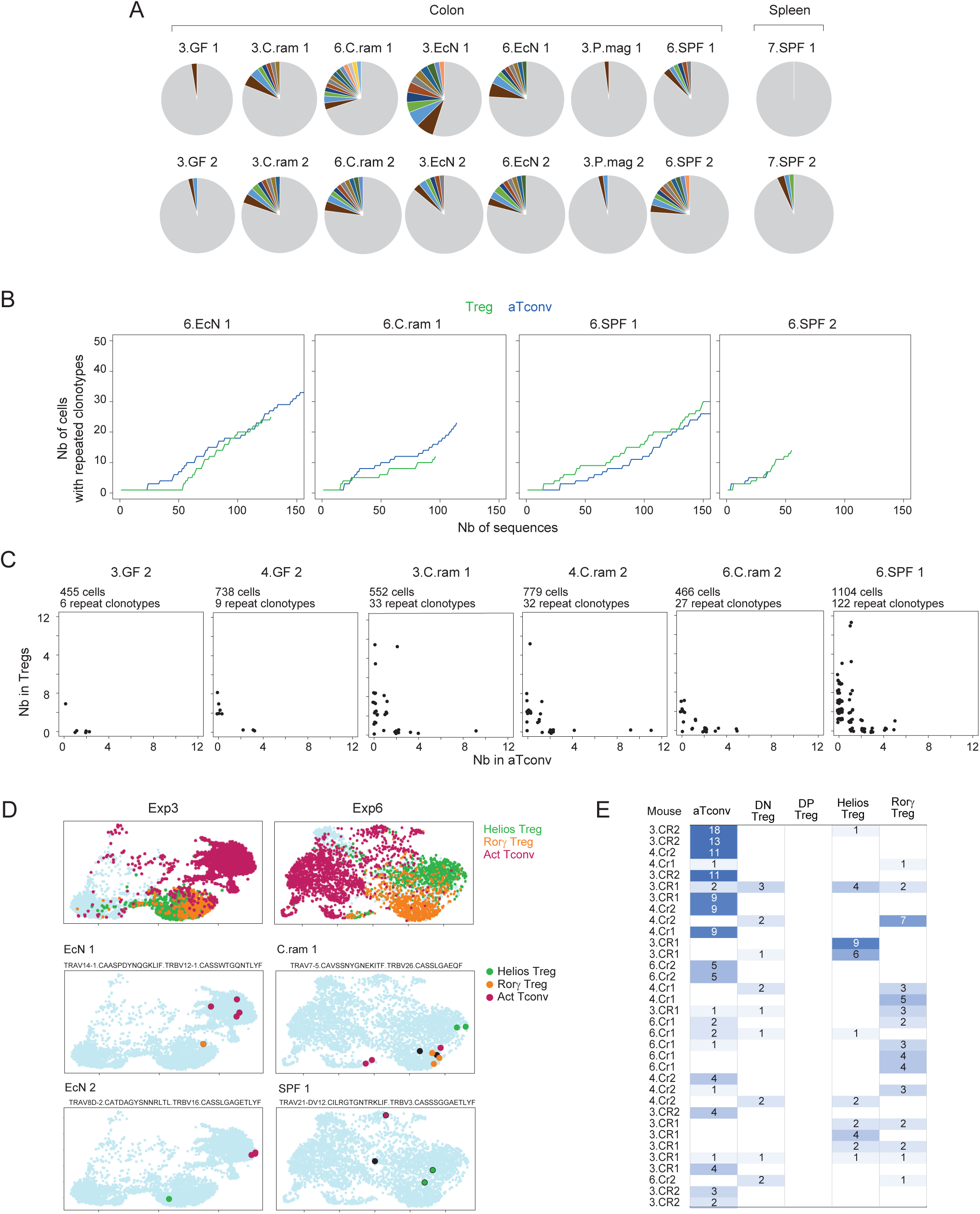
TCR clonotypes between Tconv cells and various Treg subsets. (A) Proportion of αβ TCR clonotypes in individual mice, GF, or monocolonized colon (prefix 3 and 6) and spleen (prefix 7). Nonexpanded clones in gray, colored clonotypes present in two or more cells. (B) Representative plots of number of cells with repeated clonotypes vs total number of sequences in Tregs (green) and Tconv cells (blue). (C) Representative plots of total number of repeated clonotypes in selected individual mice. (D) UMAP projection of CD4+ T cells in two separate experiments; highlights at top indicate the cell phenotype, as determined by signature gene expression; highlight on the bottom panels identify the individual cells that share the same clonotypes in the indicated mouse. (E) Summary table of amplified clonotypes in individual mice monocolonized by *C. ramosum*, and the cell-types in which they were observed.

Clonal expansions in the colon had systemic counterparts: CD4+ T cells expressing the same clonotypes were also found in splenic CD4+ T cells from the same mouse. Overall, 4.34 % of TCR clonotypes from the colon were also detected in the spleen (Table S2F). Conversely, TCRs in 3.28% of CD4+ T splenocytes had an analog in the colon, in monocolonized as well as SPF mice (Table S2G), but slightly lower in GF mice. These results suggest that gut-derived T cells make a substantial contribution to the repertoire of extra-gut lymphoid organs at steady state.

Plotting the distribution of these expanded clonotypes showed that each was preferentially found in either Tconv or Treg cells (Fig. 6C), with the strongest Tconv bias in mice monocolonized with *E. coli*, as previously reported ^49^, and in general for intestinal T cells ^4^. On the other hand, this preference was not complete, and many clonotypes were found in both Tconv and Treg of the same mouse (Fig. 6C). These occurrences did not correspond to borderline assignments of cell identity, as the cells that expressed them fell in clearly demarcated regions of the UMAP (Fig. 6D). One might expect to observe common TCRs between pTregs and expanded Tconv cells from which they had differentiated. Interestingly, however, annotation of Treg subsets using the signatures defined above revealed that there was essentially as much sharing between Tconv and Helios+ Tregs as with Rorγ+ Tregs (Table S2E, Fig, 6D). As illustrated for all the amplified clonotypes on C*. ramosum* monocolonized mice (Fig. 6E) there was also sharing between Helios+ and Rorγ+ Tregs. These results indicate that pTregs are not only Rorγ+ Tregs, and/or that the phenotypes of colonic Tregs are not cast in stone, echoing the genomic observations above.

## DISCUSSION

Using a wide and complementary array of immunologic, genomic, microbiological and functional assays, we investigated in a systematic manner the relationships between the subpopulations of colonic Treg cells that have been described in recent years, and the functional involvement of the key TFs that have been associated with them. The results point to more complex inter-relationships than simple dichotomies, as manifest in chromatin and transcriptional programs, the mode of operation of identifying TFs, or the homeostatic balance between subsets. Cell tracing via TCR barcodes also indicates that the oft-accepted tight relationship between Treg phenotypes and differentiative origin (tTreg==Helios+ Tregs, pTreg==Rorγ+ Treg) is not absolute. Instead, we propose that the setpoints resulting from microbial and other influences can modulate colonic Treg phenotypes irrespective of their locale of differentiation.

From the standpoint of the transcription factors that have been used to define these colonic Treg subpopulations, we uncovered several different modes of action. At one extreme, Gata3 seemed unnecessary to support the differentiation of any Treg subset, but influenced transcriptional activity, as manifest by chromatin accessibility profiles (the de-repression of *Il10* in *Gata3* TrKO was particularly striking, Fig. 4E). At the other extreme, both cMaf and Rorγ seemed necessary to drive the differentiation or survival of those Tregs that express them, judging from the large swings in subset frequencies in their absence (Fig. 1) with the caveat that it can be difficult to evaluate effects when the ablated TF is needed for subset identification. Helios seemed less required for the subset it marks, in light of the increase in DN Tregs (likely “Helios+ Treg wannabes”) in *Ikzf2* TrKOs, a shift that was particularly clear in the lung. Both the flow cytometry and chromatin enrichment analysis showed that factors like cMaf or Batf had broader distribution of expression and functional activity than might have been expected from prior reports.

From the standpoint of the Treg subpopulations, an important conclusion may be that they represent poles at the extreme of a spectrum, rather than cleanly demarcated entities. Several arguments bolster this view (i) The gene expression profiles, where indexing colonic Tregs in the scRNAseq results according to Helios+ and Rorγ+ Treg signatures showed a gradation of scores, going through DN Tregs, and shifted along this crescent by the presence of specific microbes (Fig. 5F) (ii) Helios+ and Rorγ+ Tregs (as defined by flow cytometry with TF markers) balance each other out in the TrKOs, with a reciprocal increase in the absence of one cell or the other. The implication is that they are competing for the same homeostatic niche, which is somewhat surprising since Rorγ+ Tregs are strongly controlled by microbes, while Helios+ Tregs are not but are strongly regulated by IL33 via ST2. Some over-arching homeostatic control must be limiting the setpoint of total colonic Tregs, irrespective of their proportion, which is tuned by microbes or maternal programming (iii) the blurry distinction between Helios+ Tregs and Rorγ+ Tregs was reinforced by the αβTCR clonotype data: several clonotypes were present in Treg cells otherwise well demarcated as Helios+ or Rorγ+ Tregs. This sharing demonstrates that these cells are directly related, either by sharing a common ancestor or by switching between phenotypes. Such sharing, in the context of a full polyclonal repertoire, aligns with earlier studies with constrained-diversity transgenic mice in which the same TCR clonotypes were found in both Helios^+^ and Rorγ^+^ Tregs ^57, 58^.

Consequently, the simple equation Helios+ Tregs == tTregs, and Rorγ+ Tregs == pTregs is no longer tenable. Cell-transfer studies with microbe-specific transgenic Tconv cells, which offer direct evidence for pTreg conversion, yielded pTregs with a predominant but not exclusive Rorγ^+^ phenotype ^8, 58, 59^. Van der Veeken et al ^13^ provided compelling evidence that the vast majority of pTregs are indeed Rorγ+ Tregs, but this was in the context of a sharp recovery from food and microbial antigen deprivation and from Treg ablation, thus in particular circumstances where pTregs are very actively generated to restore the Treg pool. In contrast, we reported earlier that the outcome of pTreg differentiation (Helios+ vs Rorγ+ Treg) depended on the type of input Tconv cell ^48^. In addition, Rorγ can be experimentally upregulated in tTregs ^60, 61^. To reconcile these observations, we thus propose the following model: pTreg generation, in the gut or elsewhere, is inherently agnostic as to Rorγ or Helios phenotype, and is driven by homeostatic factors that aim to regulate total Treg pool size. Rorγ or Helios phenotypes are instead controlled by independent factors, like microbial products, factors from particular APCs, maternally-derived setpoints, neuronal influences, etc that happen to dominate at the time of conversion. These factors may be particularly effective during the malleable stage of pTreg differentiation, but can also act on established tTregs. Which microbial structures or molecules coordinately swing this balance remains unclear ^4^.

The genomic and homeostasis results also identify an intermediate population of double-negative Tregs in the colon, not only because of their intermediate indices, but also because they selectively express a particular set of genes (Fig. 5B). We surmise that these are equivalent to the DN population described in relation to food antigens in the small intestine ^12^. Do DN Tregs represent an intermediate between Helios+ vs Rorγ+ Tregs or rather a stable and unrelated meta-state?

From a functional standpoint, the continuum perspective of colonic Treg cells does not detract from the specific functions associated with the different poles. In keeping with prior reports ^8–10, 19^, Maf+Rorγ+ Treg cells had the strongest influence on dampening inflammation in DSS colitis, possibly because of their dominant production of Il10, which would also explain their broad effects on suppressing Teff cells of different flavors, and we confirm here the inhibition of IgA production by these Treg cells. Perhaps paradoxically, *Ikzf2* TrKO mice had a less active immune response against *C. rodentium*. We suspect that this outcome may reflect the higher proportion of Rorγ+ Tregs, rather than the dearth of Helios+ Tregs, leading to “over-suppression” of the anti-microbial response. In the same vein, the higher incidence of recurring *T. muris* infection could be interpreted as an intriguing requirement for Rorγ Tregs in the anti protozoan response, or instead by over-suppression through the more abundant Helios+ Tregs. One should note that such interpretative uncertainty, when either a physiological effect on the primarily targeted cell or on a homeostatically connected cell-type can be at play, may be a more frequent issue than realized. In terms of global influence on the gut microbiota, and in contrast to ^7^, none of the TrKO mice showed marked dysbiosis, defined as severe perturbations that rearrange the balance of microbial phyla and families, or as the repeated over-representation of a specific species or genera. Alpha-diversity was comparable in all TrKOs mice, and examination of Fig. 3K and S3 shows that cage-of-origin variations are more pronounced than any mutation-specific effects. Thus, with the caveat that our experiments were only powered to identify strong dysbiosis, but not subtler effects on the microbiome network, none of the colonic Treg subpopulations seem to uniquely regulate microbial populations.

In summary, we set out to define the dynamics of colonic Treg subsets, and found that while they have functional and genomic individualities, they are interconnected in many ways, and that microbe and tissue specific cues modulate their phenotypic spectrum.

## MATERIALS AND METHODS

### Mice

C57BL/6 (B6) mice were purchased from Jackson laboratories. B6 germ-free mice were bred and maintained in our facility at Harvard Medical School. *Rorc ^fl/fl^* mice ^62^, *Ikzf2^fl/fl^* mice ^15^, *Gata3^fl/fl^* mice^63^, and *cMaf^fl/fl^* mice ^8^ were crossed to *Foxp3^cre^* mice ^64^ and were bred and maintained in our facility. All experiments were performed following guidelines listed in animal protocol IS00001257, approved by Harvard Medical School’s Institutional Animal Care and Use Committee.

### Mice treatment, infection, and colonization

For mono-colonization, GF mice were housed and orally gavaged with single bacterial species (*C. ramosum, P.magnus*, and *E. coli* Nissle) at 4 weeks of age for 2 weeks, as described ^47^. *P. magnus*, and *C. ramosum* were grown in BBL brucella blood agar plates, followed by overnight standing cultures in PYG broth under strictly anaerobic conditions (80% N_2_, 10% H_2_, 10% CO_2_) at 37°C in an anaerobic chamber. *E. coli* Nissle was grown in LB at 37°C under aerobic conditions. Stool was collected from mono colonized mice and plated at 2 weeks to verify colonization and rule out contamination from other species. For infection, eight-week-old mice were orally gavaged with 1 × 10^9^ cfu of *C. rodentium*resuspended in 100 μl PBS. *C. rodentium* was first grown on MacConkey Agar plates, followed by overnight cultures in LB with shaking at 37°C. The overnight culture was then diluted to an optical density of 0.1 followed by an additional 4 hr of growth. Bacterial density was confirmed by dilution plating. Stool was collected routinely to monitor colonization and bacterial clearance.

To check for *T. muris* colonization, stool samples (∼30mg) resuspended in 500uL PBS were assessed under the microscope at 20X magnification for the presence of *T.muris*. *T. muris* can be identified as flagellated pear shaped trophozoites (active feeding stage) about 15-30um in size. Since it is a natural infection, *T.muris* colonization is not uniform and varies between mice and housing cages, making quantification difficult. Hence, mice were assigned to a high infection group (over 100 trophozoites in a single frame at 20X under the microscope), medium infection group (between 10 and 100 trophozoites) or low infection group (fewer than 10 trophozoites) and data in Fig. 3 was derived from mice in the high infection group.

For DSS-colitis, mice were treated with 2.5% DSS for 6 days in their drinking water, followed by 4 days of recovery with normal water.

### Lymphocyte analysis and cell sorting

Intestinal tissues were measured, cleaned, and treated with RPMI containing 1 mM DTT, 20 mM EDTA and 2% FBS at 37C for 15 min to remove epithelial cells, minced and dissociated in collagenase solution (1.5mg/ml collagenase II (Gibco), 0.5mg/ml dispase and 1%FBS in RPMI) with constant stirring at 37C for 40min. Single cell suspensions were filtered and washed with 10% RPMI solution. Cells were then stained with different panels of antibodies with surface markers and intracellular markers. For cytokine analysis, cells were treated with RPMI containing 10% FBS, 30ng/ml phorbol 12-myristate 13 acetate (Sigma), 1μM Ionomycin (Sigma) in presence of GolgiStop (BD Biosciences) for 3.5 hours. For intracellular staining of cytokines and transcription factors, cells were stained for surface markers and fixed in eBioscience Fix/Perm buffer overnight, followed by permeabilization in eBioscience permeabilization buffer at room temperature for 45 min in the presence of antibodies. Cells were acquired with a BD FACSymphony and analysis was performed with FlowJo 10 software.

For scRNA-seq and T cell receptor sequencing (three independent monocolonization experiments), CD4+ T cells from distal colons of GF or monocolonized mice were sorted (DAPI CD19 CD4+ TCRβ+) after hashtagging with Biolegend TotalSeq-C reagents (10 different hashtags per experiment). All samples were pooled together in 0.04% BSA. Encapsulation was done on the 10X Chromium microfluidic instrument (10X Genomics). Libraries were prepared using Chromium Single cell 3’ reagents kit v2 according to manufacturer’s protocol. GEX, TCR and hashtag libraries were processed as described in ^49^. For scATAC-seq, colonic lamina propria cells were sorted for Tregs (DAPI-CD19-CD8-CD11b CD4+TRCRb+CD25hi) and activated Tconvs (DAPI-CD19-CD8-CD11b-CD4+TRCRb+CD25loCD44hi). Cells were hashtagged by condition at the time of staining as detailed below.

### scATAC-seq library preparation and hashtagging

Nuclei isolation, transposition, GEM generation, and library construction targeting capture of 10000 cells were carried out as detailed in the Chromium Next GEM Single Cell ATAC manual (10x Genomics) with modifications to allow sample hashtagging (see below). Libraries were pooled and sequenced on an Illumina NovaSeq 6000 to a final median depth of approximately 20-30,000 paired-end reads per cell. Sequencing data were converted to fastq files, aligned to the mm10 reference genome, and quantified per cell using Cell Ranger ATAC software (10x Genomics, v1.2).

To include multiple experimental conditions in the same scATAC run, we hashtagged cells using a modification of the ASAP-seq strategy ^65^ for low cell input primary cell samples. Before sorting, cells were hashtagged with mouse TotalSeqA DNA-barcoded hashtag antibodies at the same time as staining with fluorophore-conjugated antibodies (BioLegend). Cells were sorted into RPMI + 5% FCS in DNA Lo-Bind tubes (Eppendorf, cat # 022431021). After spinning down for 5 min at 500g in a refrigerated centrifuge at 4°C, cells were resuspended in 100 μl chilled 0.1x Omni Lysis buffer (1x Omni Lysis buffer (10mM Tris-HCl, 10mM NaCl, 3 mM MgCl_2_, 0.1% Tween-20, 0.1% NP40 substitute/IGEPAL, 0.01% Digitonin, 1% BSA in nuclease free water) diluted 1:10 in Wash/Lysis Dilution Buffer (10mM Tris-HCl, 10mM NaCl, 3 mM MgCl_2_, 1% BSA in nuclease free water), gently mixed by pipetting, and incubated on ice for 6.5 min. Following lysis, 100 μl chilled wash buffer was added and gently mixed by pipetting. Cells were spun down for 5 min at 500g at 4°C, all but 5 μl of supernatant was removed, and 45 μl of chilled 1x nuclei buffer (10x Genomics) was added without mixing. After one more centrifugation step at 500g, 4°C for 5 min, supernatant was removed, and samples were resuspended in 7ul 1x nuclei buffer for cell counting and input into transposition, barcoding, and library preparation according to the Chromium Next GEM Single Cell ATAC manual (10x Genomics).

Modifications to the original 10X protocol were made as described in the original ASAP-seq publication^65^ and as detailed at https://citeseq.files.wordpress.com/2020/09/asap_protocol_20200908.pdf. Briefly, 0.5 μl of 1uM BOA bridge oligo was spiked into the barcoding reaction. During GEM incubation, an additional 5 min incubation at 40°C was added to the beginning of the protocol. 43.5 instead of 40.5 μl of Elution Solution I was added during silane bead elution to recover 43 μl. 40 μl was used for SPRI clean up as indicated in the protocol, while 3 μl was set aside. During SPRI cleanup, the supernatant was saved. The bead bound fraction was processed as in the protocol, while for the supernatant fraction, 32 μl SPRI was added for 5 min. Beads were collected on a magnet, washed twice with 80% ethanol, and eluted in 42 μl EB. This 42 μl was combined with the 3 μl set aside from the previous step as input into the HTO indexing reaction. HTO Indexing PCR was run with partial P5 and indexed Rpxx primers (https://citeseq.files.wordpress.com/2020/09/asap_protocol_20200908.pdf) as: 95°C 3 min, 12-14 cycles of (95°C 20 sec, 60°C 30 sec, 72°C 20 sec), 72°C 5 min. The PCR product was cleaned up with 1.6X SPRI purification for quantification and sequencing alongside ATAC libraries.

scATACseq data analysis was performed using Signac v1.4^66^. For quality control, only cells with at least 1x10^3^ fragments per cell (depending on sequencing depth of experiment), greater than 25 percent reads in peaks, TSS enrichment score greater than 2, nucleosome signal less than 10, and ratio of blacklist region reads less than 0.05 were retained for further analysis. Putative doublets identified by ArchR v1.0.1^42^ and non-Treg, non-Tconv contaminant cells were also removed. We used the latent semantic indexing approach as previously described^67, 68^. Binarized count matrices were normalized using the term frequency-inverse document frequency (TF-IDF) transformation and reduced in dimensionality by singular value decomposition (SVD). As the first component was highly correlated with sequencing depth, SVD components 2-30 were used to generate a shared nearest neighbor (SNN) graph for clustering and as input into UMAP^69^ with cosine distance metric for visualization.

*Hashtag counts + assignments*. Hashtag processing followed the original recommendations of the ASAP-seq paper^65^, using asap_to_kite (https://github.com/caleblareau/asap_to_kite) to process FASTQs files for downstream quantification by the bustools and kite workflows^70, 71^. We used the HTODemux()^72^ function in the Seurat package (v4.0.2)^73^ to remove doublets and call hashtag identities. Because of some variation in absolute hashtag numbers leading to inconsistent hashtag assignment using HTODemux(), this procedure was supplemented with classification of cells by clustering based on normalized hashtag counts and visualization using PCA or UMAP of hashtag counts.

#### Peak Sets

To enable comparisons across conditions and datasets, we used a common set of Treg specific open chromatin regions, defined previously ^14^ by supplementing pan-immunocyte OCRs from the ImmGen consortium^44^ with additional peaks from Treg spleen and colon scATAC-seq data in ^14^. To arrive at the final peak set, we also called peaks on the data generated in this study, using Archr v1.0.1^42^ and MACS2 v2.2.7.1^74^ to call fixed-width peaks of 250bp within cell clusters. Any new OCRs not overlapping with the previous peak sets were added to the final merged peak set used for downstream analysis.

#### Motif Accessibility Analysis

Bias-corrected relative motif accessibility was calculated using chromVAR^43^. We used motifmatchr (https://github.com/GreenleafLab/motifmatchr) to scan OCRs in our refence set from the curated set of mouse motif PWMs from the Buenrostro lab (https://github.com/buenrostrolab/chromVARmotifs/tree/master/data/mouse_pwms_v2.rda).

#### Gene Scores

Gene scores were calculated with Archr v1.0.1, using an exponentially weighted function that accounts for the activity of distal OCRs in a distance-dependent manner ^42^ and provides an approximate proxy for gene expression.

#### Pseudobulk Track Visualization

To visualize pseudobulk profiles, BAM files containing reads for each group of cells were extracted using Sinto (https://github.com/timoast/sinto), shifted to account for Tn5 cut-sites, and converted to bigwigs using deeptools^75^ for display in the Integrative Genomics Viewer^76^.

#### OCR Signature Relative Accessibility

Per-cell relative accessibility of OCR sets, including signature OCRs, TF binding sites, etc, was calculated using the chromVAR computeDeviations() function^43^. Helios+ vs Rorγ+ accessibility signatures were retrieved from ^14^ and Rorγ ChIP-seq bound sites were from the ImmGen consortium ^44^.

### Single-cell RNAseq processing for TCR and gene expression analysis

For GEX, data were analyzed in R using the Seurat package ^77^. HTOs were assigned to cells using the HTODemux function, and doublets were eliminated from analysis. Cells with less than 700 UMIs or 500 genes and more than 2,500 UMIs, 10,000 genes and 5% of reads mapped to mitochondrial genes were also excluded from the analysis. Dimensionality reduction, visualization and clustering analysis were performed in Seurat using the NormalizeData, ScaleData, FindVariableGenes, RunPCA, FindNeighbours (dims=1:30), RunUMAP (dims=1:30) and FindClusters functions. Cluster identity was determined based on expression of key marker genes. Assignment of Tregs, Rorγ+ Tregs and Helios Tregs were based on signatures obtained from Pratama et al ^48^: (Activated CD4 T cells (**ActT**): *Cd44, Icos, Gata3, Pdcd1, Rorc, Tbx21* **Tregs**: *Foxp3, Il2ra, Il10, Lag3, Tigit* **Rorg+ Tregs:** *Cd200r1, Cd83, Dgat2, Epas1, Fam46a, Gas2l3, Ikzf2, Il9r, Naip5, Nrp1, Ppp2r3a, Swap70* **Helios+ Tregs:** *Ccr1, Ccr2, Ccr5, Ccr9, Clic4, F2rl2, Gpr155, Havcr2, Itgb5, Marcks, Matn2, Nr1d1, Prg4, Rorc*). For 2D representation of Tregs, scores were obtained based on the gene signatures listed using AddModuleScore function and then plotted using ggplot2.

For TCR, TCRα and TCRβ data were generated using the CellRanger v6.1.0 VDJ pipeline. Output files included a CSV file labelled as filtered_contig_annotations.csv, which contains clonotype information for barcoded cells, and another labelled as all_contig.fasta, which contains TCRα and TCRβ sequences on a per-barcode basis. The filtered_contig_annotations.csv file contains CellRanger-predicted V, D, J, and C genes for alpha and beta chains, as well as corresponding CDR1-3 and FWR1-4 sequences. The corresponding entries in the all_contig.fasta file contains full V through joining region sequences as well as the beginning of the constant region. These clonotype information along with sequence information was analyzed and stored using Seurat so that each cell would have all the relevant TCR information for both the alpha and beta chains. Clonotypes were determined by the combination of TRAV, CDR3a, TRBV and CDR3b regions and sequences, at the protein level. In every instance in this study, clonotypes found to be repeated in one mouse invariably corresponded to the exact same DNA nucleotide sequence for both alpha and beta chains, hence likely resulted from clonal expansion (occurrences of identical clonotypes in different mice showed divergence at the nucleotide level). To estimate the frequency of repeated clonotypes, without confounding by the variable depth of sequencing for different samples, random sets of 80 clonotypes were drawn iteratively from each sample (100 iterations) and the mean frequency of clonotypes observed more than once was averaged between the draws.

For rearranged TR gene junction analysis, we first identified the extent of TRJ and TRV trimming, or the eventual inclusion of P nucleotides, by alignment with germline reference sequences obtained from the IMGT website (https://www.imgt.org/). P nucleotides were scored as germline sequence. For TRA genes, the remaining nucleotides were classified as N additions. For TRB genes, these nucleotides were analyzed for the compatibility with a TRBD origin. The un-assigned nucleotides were classified as N additions.

### Bacterial population profiling

Every TrKO and its paired control littermate (sex-matched) were selected from experimental batches spanning 3 years. DNA was isolated from stool samples using phenol/chloroform and the QIAquick PCR purification kit (Qiagen). For 16S rDNA profiling, the V4 region of 16S rRNA gene was amplified with primers 515F and 806R ^78^, and ∼390-bp amplicons were purified and then subjected to multiplex sequencing (Illumina MiSeq, 251 nt x 2 pair-end reads with 12 nt index reads, all primer sequences listed in Table S2). Raw sequencing data were processed with the QIIME2 suite ^79^. Biefly, raw sequencing data were imported to QIIME2 and demultiplexed, then DADA2 were used for sequence quality control and feature table construction. The feature table were further used for taxonomic analysis and differential abundance testing: The phylum and family taxonomy plots were generated by the function “qiime taxa barplot” and the alpha rarefaction curves were generated by the function “qiime diversity alpha-rarefaction”. The features characterizing the differences between conditions were identified by LEfSe ^80^ (Fig. S3C)

## Quantification and statistical analysis

Data are presented as mean ± SD. Unless stated otherwise, significance was assessed by Student’s t-test or Mann-Whitney using R or GraphPad Prism 8.0.

## Data availability

16S raw data and single cell sequencing data are available in NCBI with accession number XXXXX and GSE213200.

## ACKNOWLEDGEMENTS

We thank Drs X. Chi, A Gelineau, Y. Zhu, J. Léon, M. Wu for insightful discussions, K. Hattori, M. Sleeper and J, Nelson, Liang Yang B. Vijaykumar for help with mice, cell sorting and computational analysis, C. Laplace for figures. Supported by grants from the NIH (AI125603, AI150686). DR was supported by Damon Runyon Cancer Research Foundation (DRG 2300-17, National Mah Jongg League), K.C was supported by NIGMS grants T32GM007753 and T32GM144273 and a Harvard Stem Cell Institute MD/PhD Training Fellowship.

## SUPPLEMENTARY FIGURE LEGENDS

**Supplementary Figure 1:**
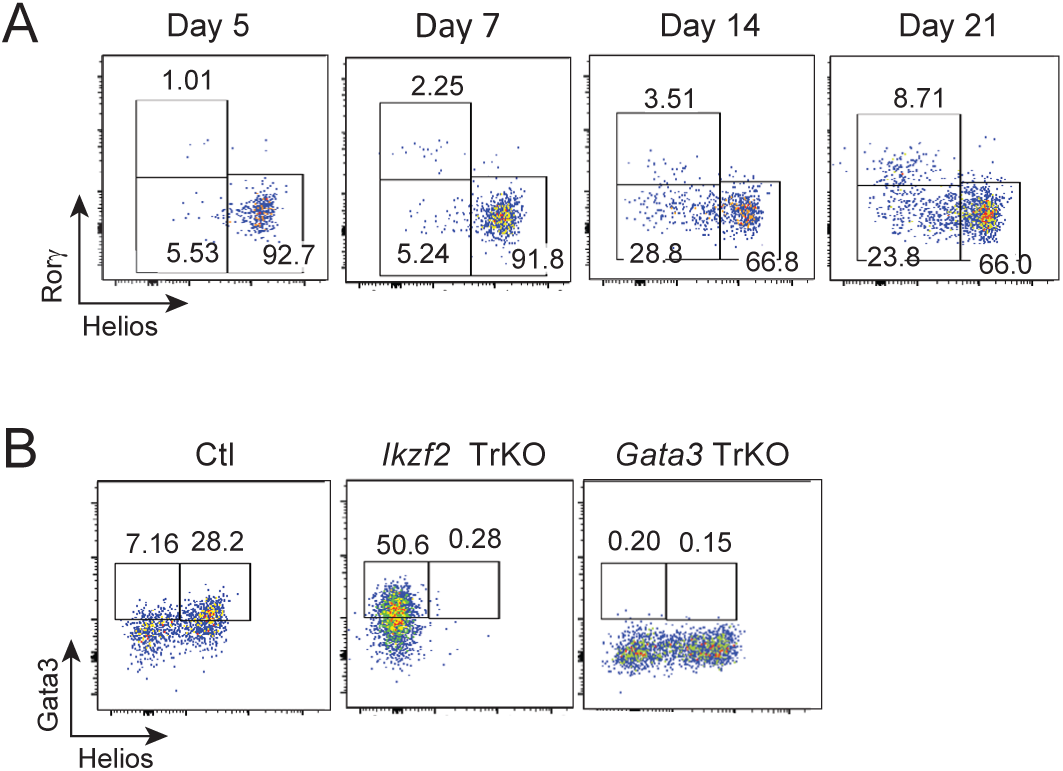
Treg subset dynamics in early life at steady-state and in the lung in TrKO mice. (A) Representative flow cytometry plots of intestinal Tregs across different time points from early life to weaning gated on TCRβ+ CD4+ Foxp3+ Tregs. (B) Representative flow cytometry plots of dominant lung Treg subsets (Helios+ Tregs and Gata3+ Tregs) in the absence of key transcription factors.

**Supplementary Figure 2:**
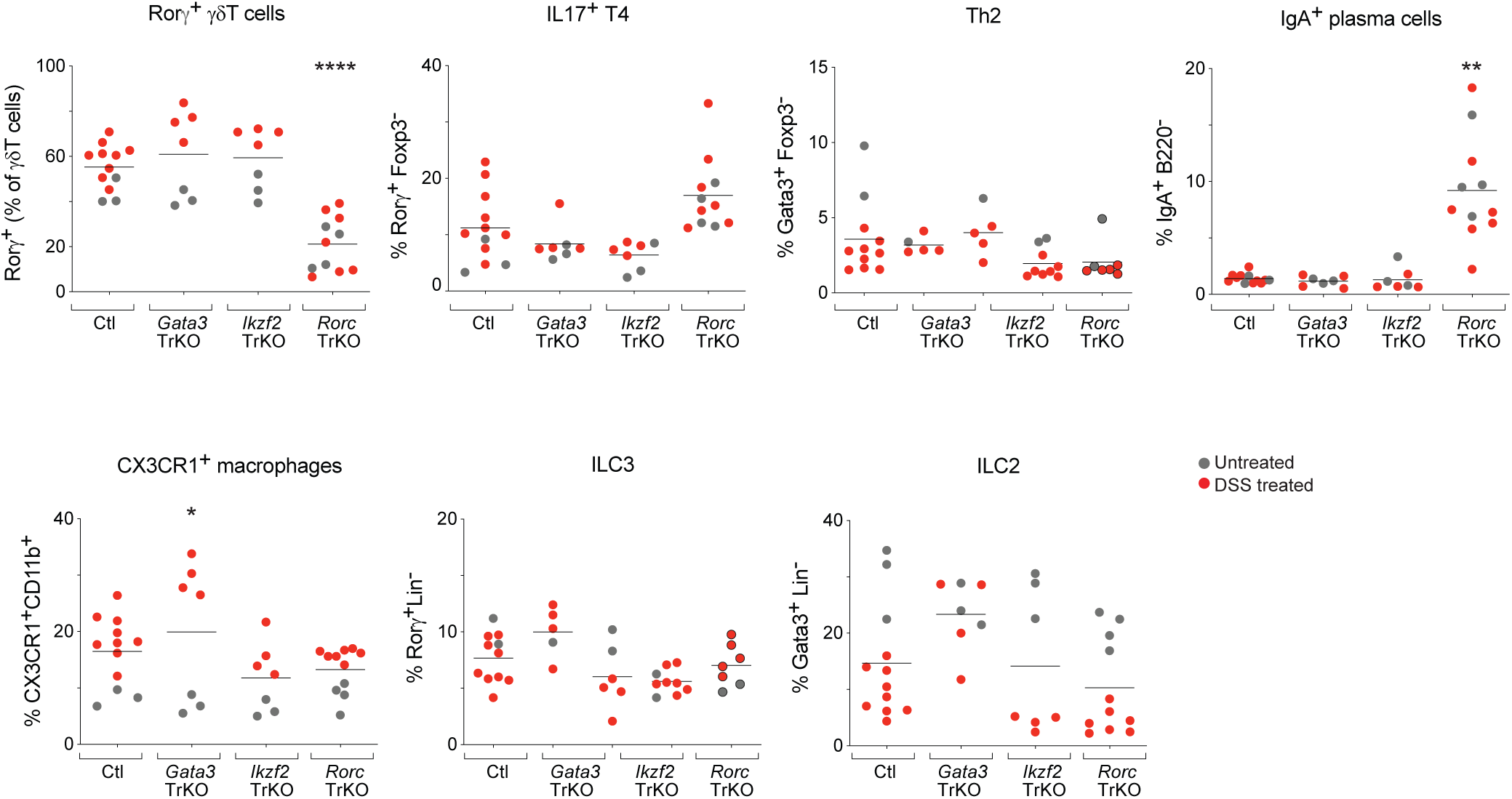
Regulation of colonic immune cells in untreated and DSS-treated TrKO mice. (A) Quantification of selected colonic immune cell types in different TrKO and pooled littermate controls. **p<0.01 and ****p<0.0001 by unpaired t-test.

**Supplementary Figure 3:**
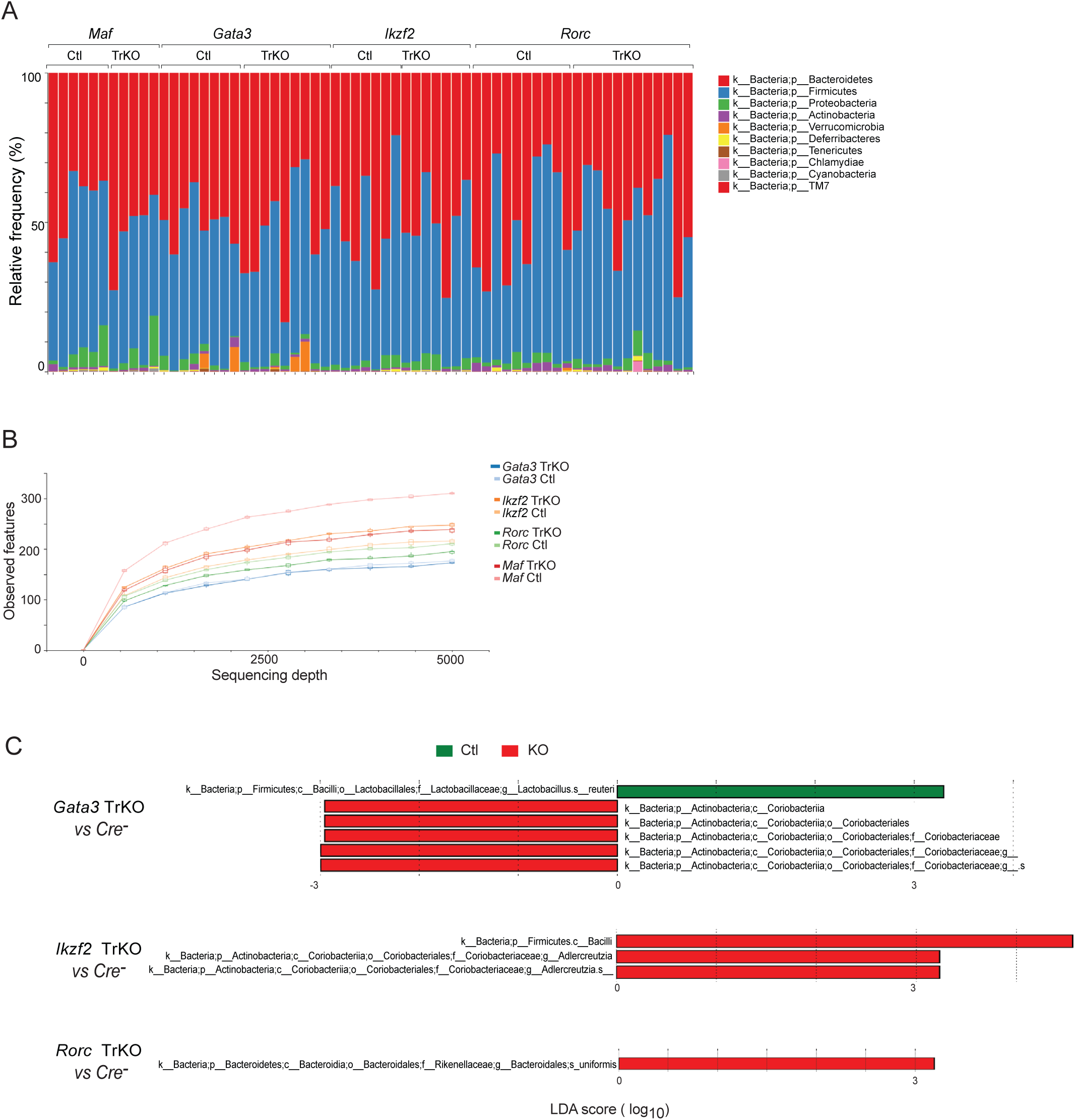
Analysis of microbial populations in stool reveals no major differences between TrKO mice and controls. (A) Relative abundance of bacterial phyla in stool of TrKO mice of the indicated groups generated by 16S rRNA sequencing. (B) Alpha diversity rarefaction plot of microbial populations in TrKO mice and control littermates. (C) LEfSe (Linear discriminant analysis Effect Size) analysis of microbial communities in each TrKO compared to paired littermate control

**Supplementary Figure 4:**
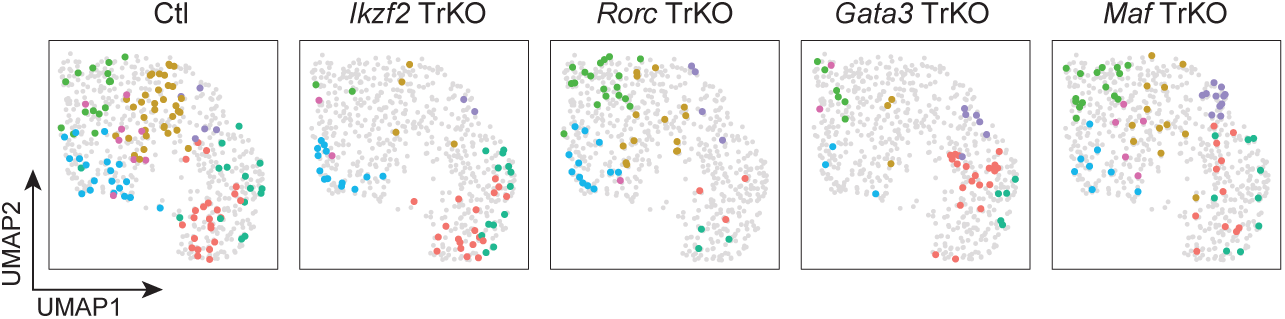
Genomic identities of intestinal Tregs in TrKO mice. (A) UMAP of scATAC-seq data of intestinal Tregs as in fig. 4A split by each TrKO and control (identified by individual hashtags).

**Supplementary Figure 5:**
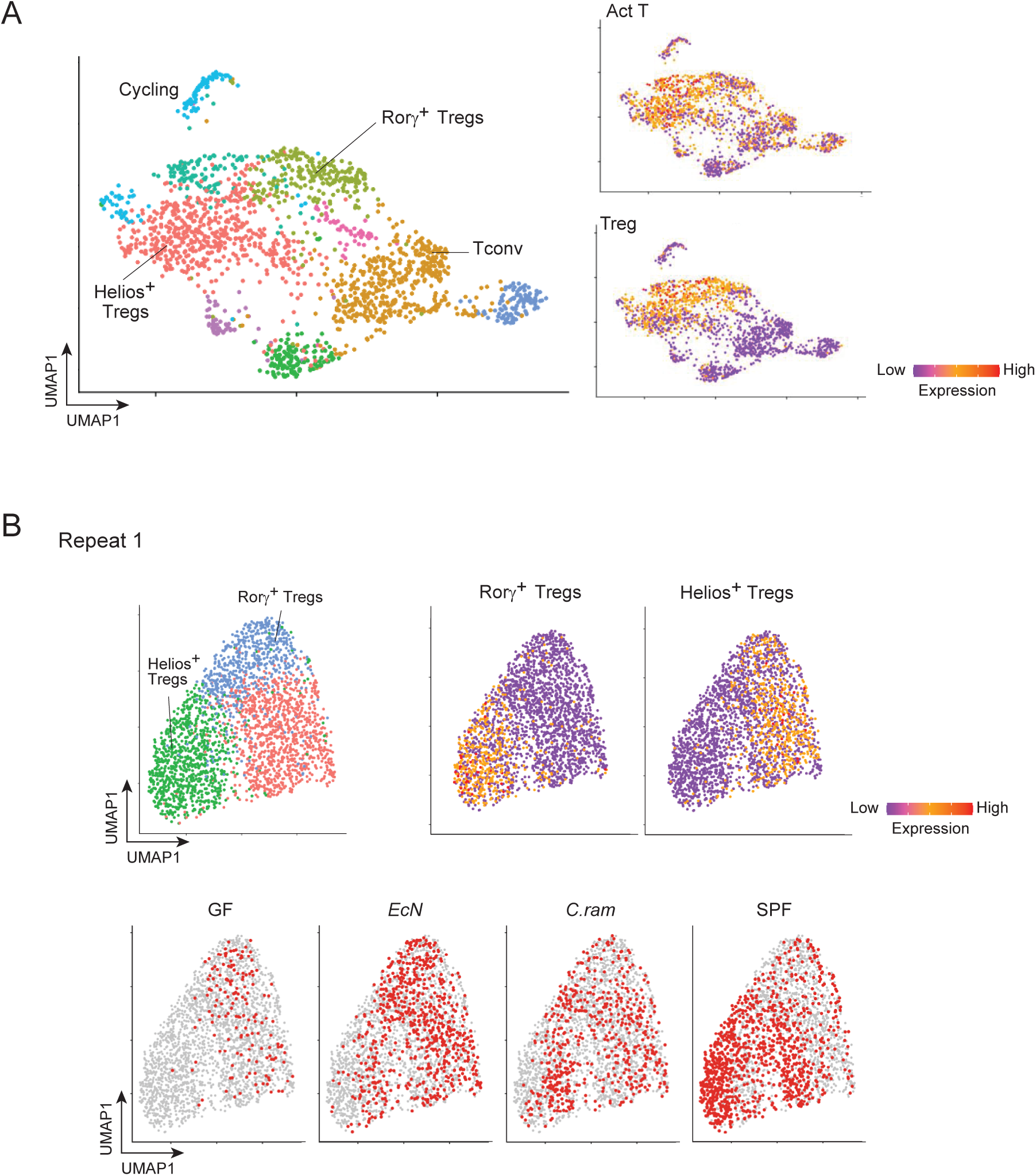
Helios+ Tregs and Rorγ+ Tregs have unique transcriptional signatures that are conserved across responses to individual microbes. (A) UMAP projection of colonic CD4 T cells clusters from a representative scRNAseq analysis (left), color-coded for expression of activated CD4 T cell and Foxp3+ Treg gene signatures (right) (genes in Materials and Methods) (B) UMAP projection of colonic Treg cell clusters from a representative scRNAseq analysis of colonic Tregs (repeat of experiment in fig. 5), color-coded for expression of RORγ+ and Helios+ Treg gene signatures (genes in Materials and Methods) (top). UMAP projection of colonic Treg cells split between hashtags from different mice. Each hashtag represents an individual mouse of the listed condition.

**Supplementary Figure 6:**
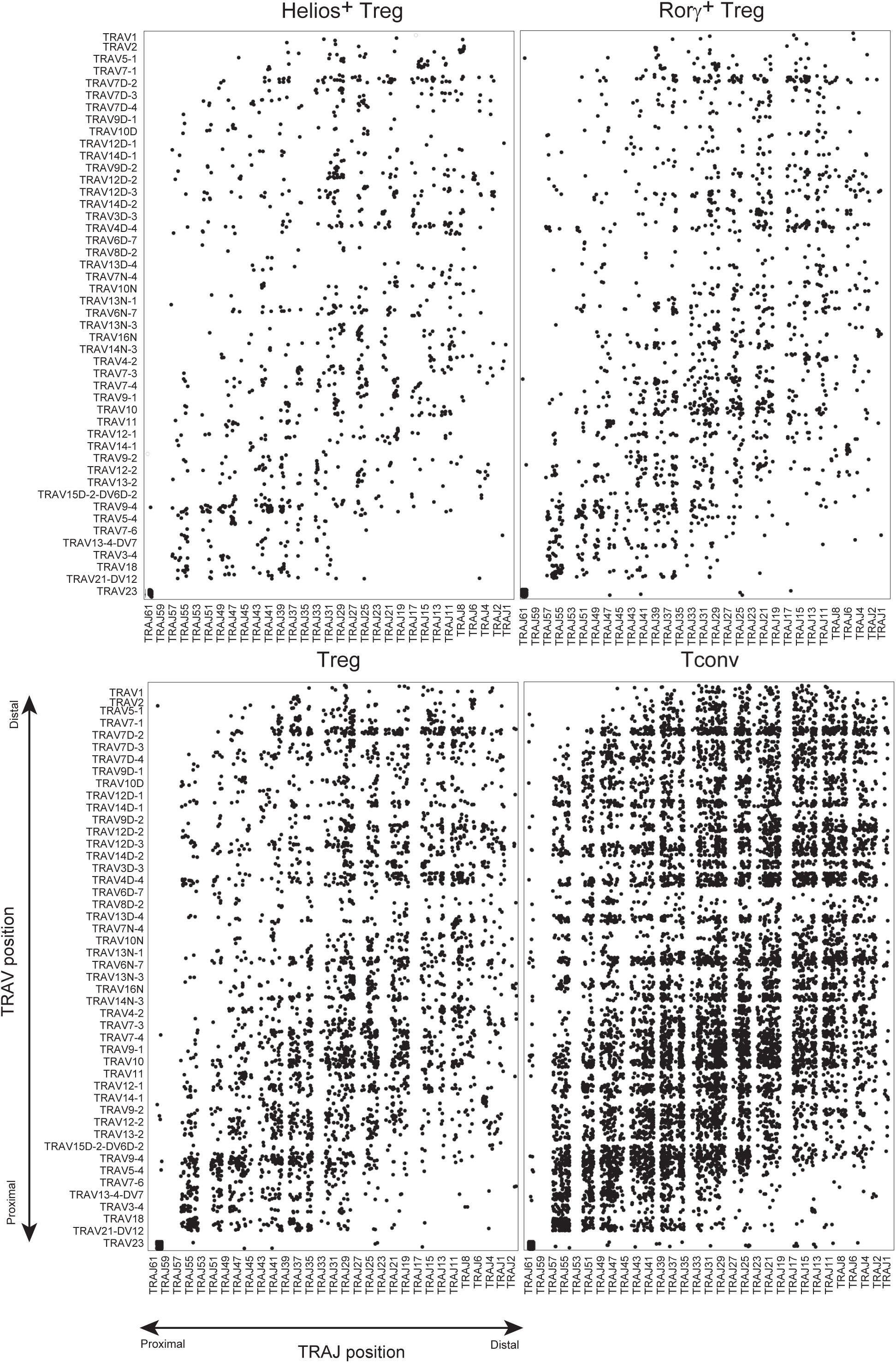
Distribution of Va and Ja regions of *Tcra* sequences from Treg subsets are comparable to Tconv cells.

**Supplementary Figure 7:**
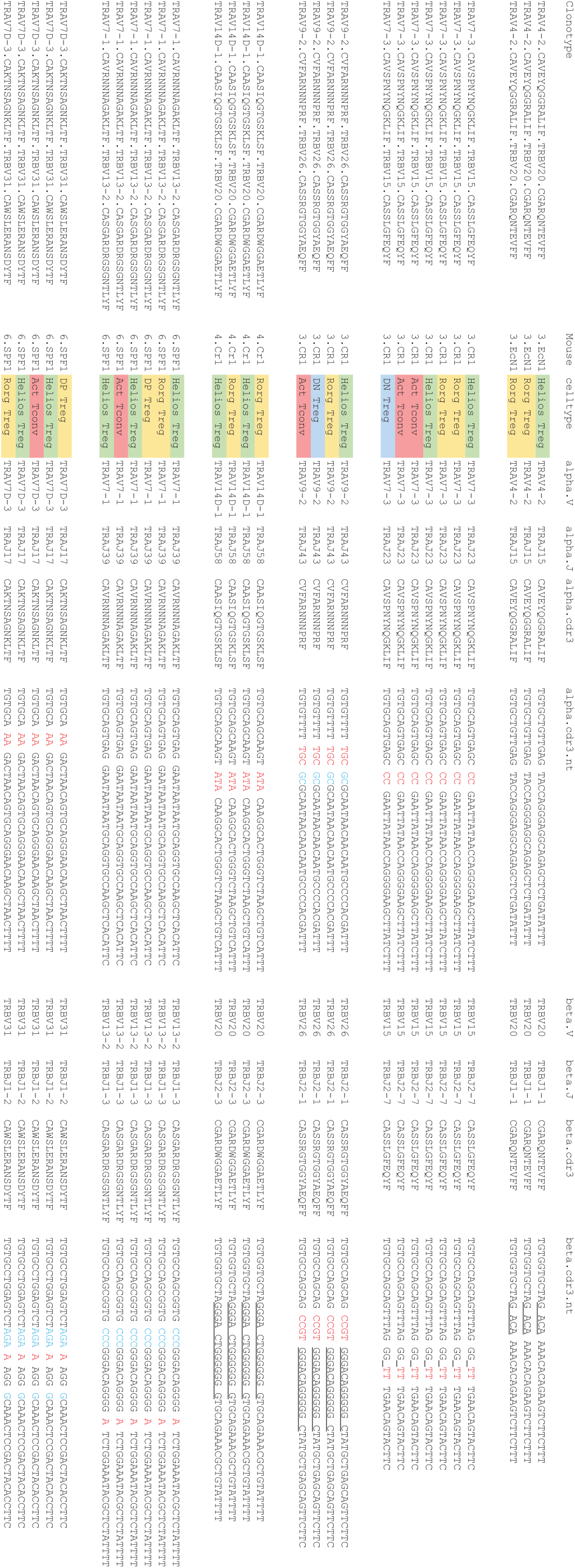
Examples of repeated TCR clonotypes repeated within one mouse. Sequences present in individual cells, parsed into TRA and TRB components. Red nucleotides: N additions; blue: P nucleotides; underlined: putative homology-mediated recombination events.

**Supplementary Figure 8:**
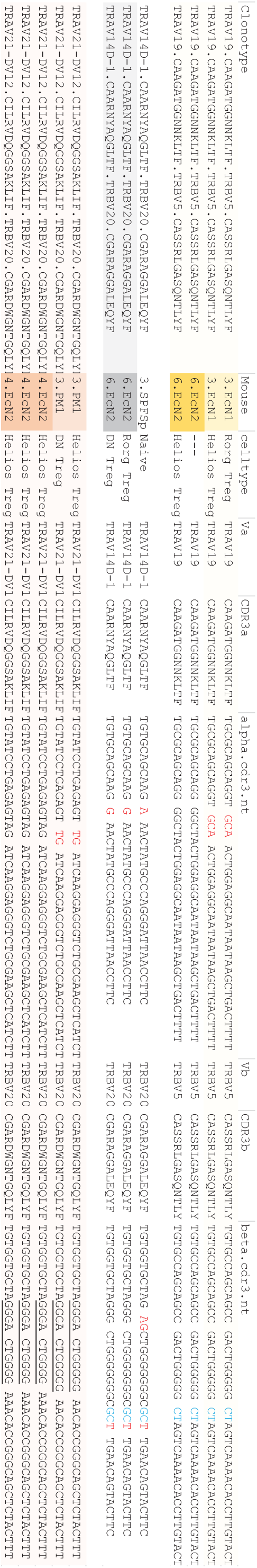
Examples of identical CDR3 regions in repeated TCR clonotypes across different mice. Sequences present in individual cells from the same or different mice (color-coded by mouse). Sequence parsing as in Fig. S7.

